# RNA-Dependent RNA Polymerase From SARS-CoV-2. Mechanism Of Reaction And Inhibition By Remdesivir

**DOI:** 10.1101/2020.06.21.163592

**Authors:** Juan Aranda, Modesto Orozco

**Affiliations:** Institute for Research in Biomedicine (IRB Barcelona), The Barcelona Institute of Science and Technology, Baldiri Reixac 10, 08028 Barcelona, Spain; Departament de Bioquímica i Biomedicine, Universitat de Barcelona, Universitat de Barcelona, Avinguda Diagonal 645, 08028 Barcelona, Spain

## Abstract

We combine sequence analysis, molecular dynamics and hybrid quantum mechanics/molecular mechanics simulations to obtain the first description of the mechanism of reaction of SARS-CoV-2 RNA-dependent RNA polymerase (RdRp) and of the inhibition of the enzyme by Remdesivir. Despite its evolutionary youth, the enzyme is highly optimized to have good fidelity in nucleotide incorporation and a good catalytic efficiency. Our simulations strongly suggest that Remdesivir triphosphate (the active form of drug) is incorporated into the nascent RNA replacing ATP, leading to a duplex RNA which is structurally very similar to an unmodified one. We did not detect any reason to explain the inhibitory activity of Remdesivir at the active site. Displacement of the nascent Remdesivir-containing RNA duplex along the exit channel of the enzyme can occur without evident steric clashes which would justify delayed inhibition. However, after the incorporation of three more nucleotides we found a hydrated Serine which is placed in a perfect arrangement to react through a Pinner’s reaction with the nitrile group of Remdesivir. Kinetic barriers for crosslinking and polymerization are similar suggesting a competition between polymerization and inhibition. Analysis of SARS-CoV-2 mutational landscape and structural analysis of polymerases across different species support the proposed mechanism and suggest that virus has not explored yet resistance to Remdesivir inhibition.

## INTRODUCTION

SARS-CoV-2 (Covid19) emerged in China in December 2019 and rapidly spread over the world, causing a world-wide health threat. At the beginning of June 2020 Covid19 pandemic has caused around 400,000 fatalities and at least 6 million people have been infected, producing an unprecedent stress to the health-care systems.^1^ From a phylogenetic point of view SARS-CoV-2 belongs to the β- genus of the coronavirus family which includes other highly infective pathogens such as the Middle East respiratory syndrome CoV (MERS-CoV) or the severe acute respiratory syndrome CoV (SARS-CoV).^2^ SARS-CoV-2 has a large (30 kb of positive RNA) genome, which forces it to a high fidelity in replication and accordingly to a relatively low mutational rate.^3, 4^ A high-fidelity RNA-dependent RNA polymerase (RdRp) and a proof-reading exonuclease^5^ are the crucial elements for maintaining the stability of the viral genome.

RdRp is the core of the replication machinery of the virus, and the largest protein in the viral genome (932 residues). It binds to nsp7 and nsp8 to form an active complex,^6^ that using as template the sense RNA generates a negative copy, which in a second cycle is used to generate new copies of genomic and sub-genomic RNAs. At least two other proteins are involved in the replication process: a 601-residue helicase and a 527-residue proof-reading exonuclease.^7^ A simple BLAST^8^ query shows that SARS-CoV-2 RdRp is highly conserved within the Coronavirus family, but no homologs are found out of it. This would suggest that we are in front of a quite new protein, which, in principle, did not have enough evolutionary time to optimize its function. Unfortunately, the extremely high infectivity of the virus suggests that despite its youth, RdRp is very effective. How the enzyme achieves a good specificity and an excellent catalytic power is still unclear, generating an important gap in our knowledge on SARS-CoV-2 replicative cycle.

Due to its central role in the viral infective cycle, RNA polymerases are a major target for fighting RNA-viruses.^6, 9^ As today, the only FDA-approved drug for treatment of SARS-CoV-2 infection^10^ is a RdRp inhibitor Remdesivir (R), a C-nucleoside (see Supplementary Fig. 1) which was in clinical trials for the treatment of Ebola,^11^ a negative single strand RNA virus very distant from SARS-CoV-2, but whose replication is also dependent on the action of a RdRp. The mechanism of inhibition of RdRp by Remdesivir is mostly unknown and conflicting hypothesis have emerged the last months. This lack of knowledge dramatically hampers our ability to develop new and more active derivatives.

**Fig. 1.**
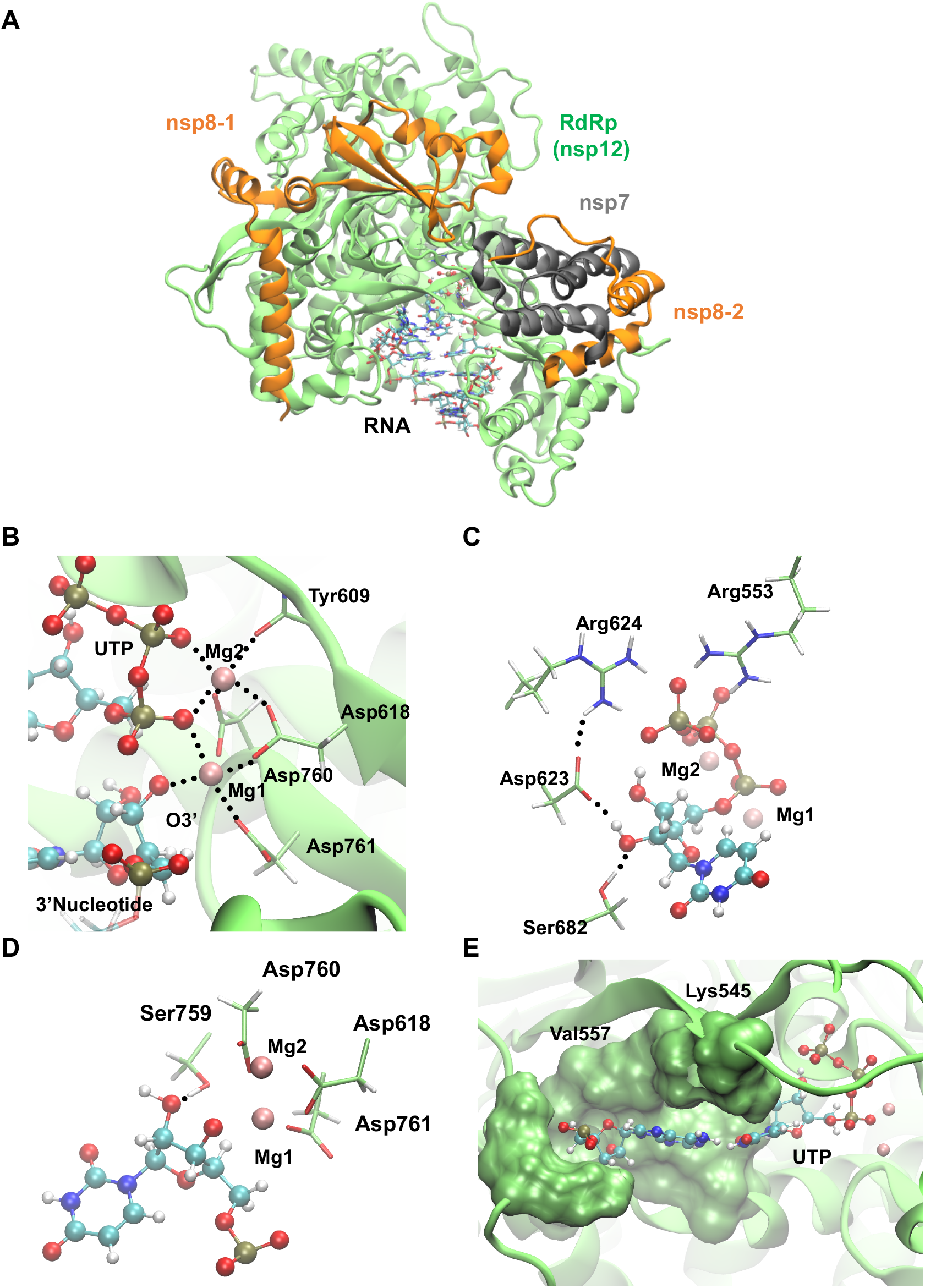
Active site of RdRp of SARS-CoV2 is an efficient polymerase. **A** Replication complex of SARS-CoV-2 used in this study is formed by the nsp12 RdRp enzyme, the nsp8 and nsp7 cofactors and RNA template and nascent strands in solution. **B** Active site insight showing the well-defined two metal ions coordination sphere. **C** Substrate recognition by active site residues of RdRp. **D** Interactions between the 3’ terminal nucleotide and RdRp. **E** RdRp pocket enables base specificity between the incoming nucleotide and the template.

We present here a comprehensive theoretical study on the mechanism of action of SARS-CoV-2 RdRp. MD simulations show that despite its youth the enzyme has a discriminant active site which should guarantee high replication fidelity. MD and QM/MM simulations demonstrate that the enzyme follows a canonical 2-ions reaction mechanism with a catalytic efficiency similar to that of the highly evolved eukaryotic polymerases. The same type of calculations demonstrates that Remdesivir triphosphate (RTP; the expected bioactive form of Remdesivir) can be recognized and incorporated into a nascent RNA with an efficiency only slightly lower than natural nucleotides, i.e. R is not an inhibitor of nucleotide incorporation. Extended MD simulations failed to detect any dramatically distortion in the RNA duplex due to the presence of Remdesivir which would justify inhibitory properties. Furthermore, no steric clashes were detected when the nascent RNA duplex was displaced along the exit channel (see below), which argue against the hypothesis that steric clashes could explain a delayed inhibition. However, the same analysis suggests that when three more nucleotides are incorporated to the RNA after Remdesivir, the C1’ nitrile group of Remdesivir reaches the proximity to Ser_861_ favoring an unexpected crosslinking reaction. Activation barriers of polymerization and crosslinking are comparable, raising an interesting scenario of competition between polymerization and inhibition which can explain the unique inhibitory properties of Remdesivir. In summary, our results provide then, not only the first atomistic description of the mechanism of action or SARS-CoV-2 RNA polymerase, but also a plausible explanation for the inhibitory mechanism of Remdesivir.

## METHODS

### RdRp complex set-up

The cryo-EM structure of the RdRp (nsp12) of SARS-CoV-2 complexed to nsp8 and nsp7^12^ cofactors was used as our starting model. We aligned it with the RdRp from Hepatitis-C virus X-ray structure^13^ from which RNA, the two catalytic metal ions and the diphosphate group of an NTP analogue were extracted (see Supplementary Methods for details). During the preparation of this manuscript a cryo-EM structure of RdRp in complex with a full RNA strand appeared^14^, confirming the accuracy of our model (see Supplementary Fig. 2). Afterwards, we built a UTP or RTP molecule inside the active site in the expected orientation required for incorporation into the nascent RNA strand. The system was solvated and neutralized prior to optimization, thermalization (T=298K) and equilibration (see Supplementary Methods for details). The final equilibrated structure was the starting point for further MD simulations. The progression of the polymerization process was simulated by adding additional base pair steps to the RNA duplex, moving the r(R·U) pair from i+1 to i+4 position keeping constant the reactive alignments at the i-position (see Supplementary Fig. 3). These structures allowed us to trace the sliding of the nascent RNA duplex along the exit path and check for potential reasons for the R-induced delayed termination of the polymerization reaction.

**Fig. 2.**
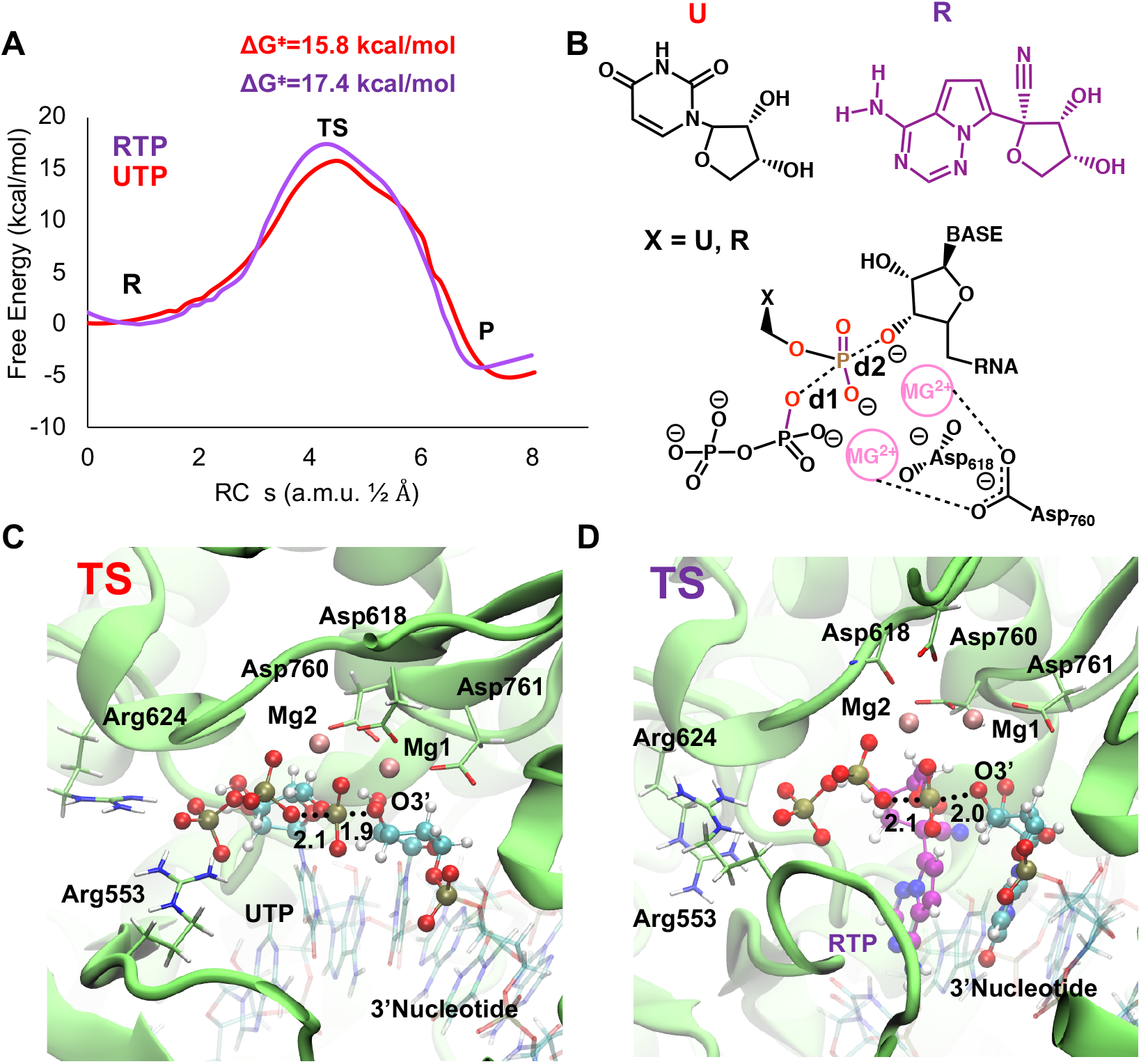
Viral RNA elongation inside RdRp of SARS-CoV-2. **A** Free energy profiles for the incorporation of a U or a R to a nascent viral RNA strand. The reaction consists on a nucleophilic attack of the O3’ of terminal nucleotide to the Pα of the triphosphate nucleotide. **B** Schemes representing the transition states found for UTP or RTP substrates. Distances included in the definition of the Reaction Coordinate (s) are shown. **C** Active site insight of the TS found when UTP is the substrate for the elongation reaction. The phosphoryl group is half-way to be transferred. Mg1 activates the O3’ towards the nucleophilic attack and stabilizes the negatively charged TS. Mg2 stabilizes the charged TS, and the newborn negatively charged PPi molecule. Distances involved in the reaction are shown as dotted lines with its average value in Å. **D** Active site insight of the TS found when RTP is incorporated (shown as ball and sticks with its C atoms shown in purple).

**Fig. 3.**
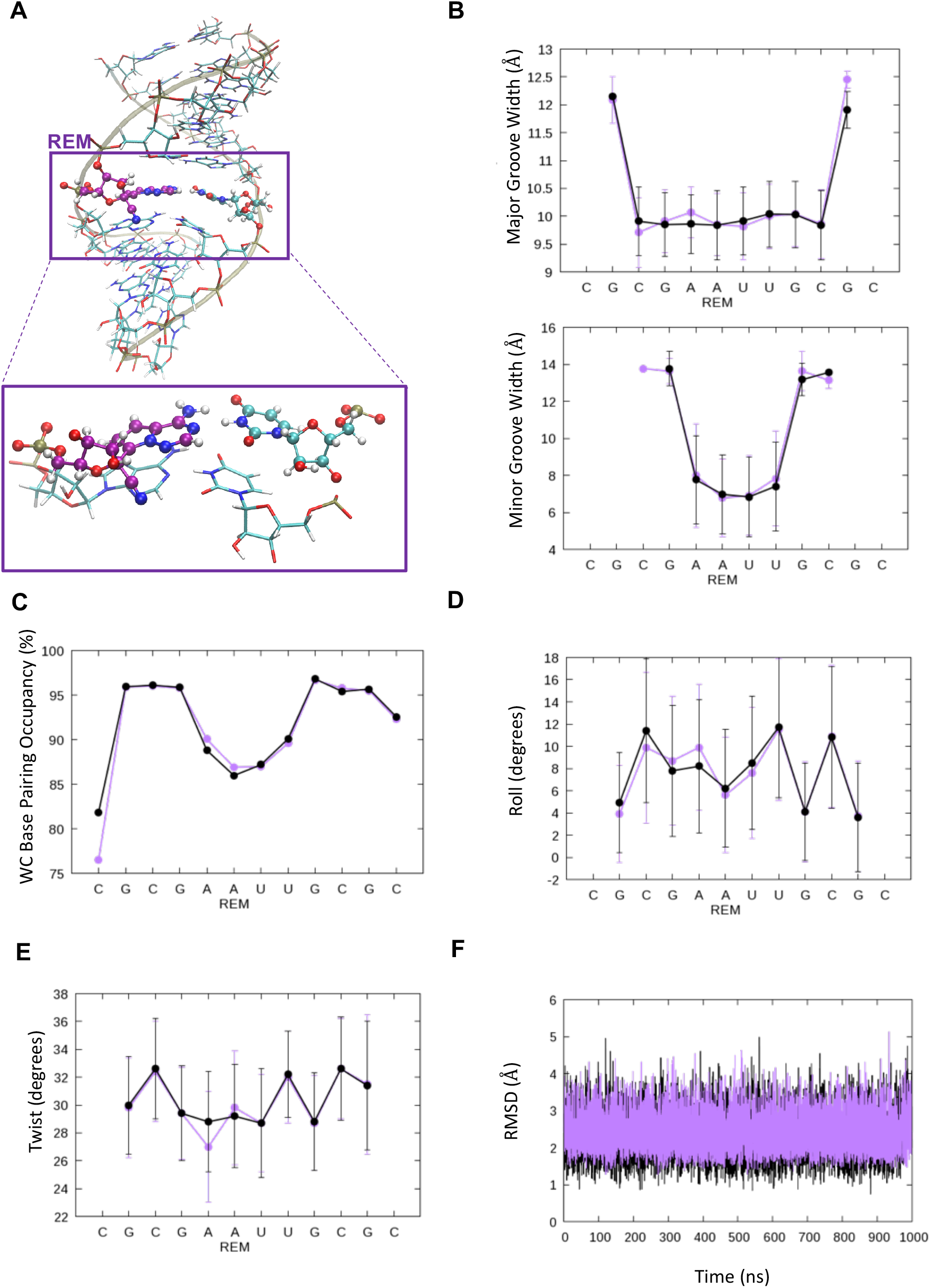
Remdesivir does not distort RNA structure. **A** Overall view of an RNA duplex containing a Remdesivir nucleotide (shown as ball and sticks with its C atoms shown in purple). An insight is shown. Major and minor groove width **B**, WC base pairing **C**, Roll **D**, Twist **E**, and RMSD of the double helix **F** are not affected when Remdesivir is present (purple) in respect to control sequence (black).

### MD simulations on RdRp

Classical trajectories were used to refine and check the stability of complexes prior to running QM/MM simulations, as well as to determine the impact of R-incorporation in RNA. Minimization, thermalization and equilibration were performed using standard procedures as described in Supplementary Methods. Production simulations were carried out using the AMBER 18 program^15^ and state-of-the art conditions for a total time of at least 0.5 μs. Water molecules were described through the TIP3P^16^ model, parameters of Magnesium ions were taken from Allner et al.,^17^ Carlson et al.^18^ parameters were used for triphosphate groups, PARMBSC0-chiOL3 for RNA.^19–22^ and ff14SB^23^ for the proteins. Parameters and charges of UTP, RTP, Remdesivir nucleotide and 3’ terminal U and R were derived to be compatible with the force fields making use of the RED server.^24^ Additional details of the simulation setups can be found in Supplementary Methods.

### QM/MM systems

Two families of systems were generated to explore enzyme reactivity. The first was created to follow nucleotide incorporation, while the second was intended to study a potential crosslinking between Ser_861_ and the nitrile group of RNA-incorporated Remdesivir. QM subsystems used in reactivity calculations are shown in Supplementary Fig. 4. The QM part was described at the B3LYP/6-31G** level of theory, using the link-atom to join QM and MM regions.^15^ The hybrid QM/MM models were built using randomly selected snapshots obtained in the last ns of unrestrained MD simulations, which were then minimized and re-equilibrated at the B3LYP/6-31G**/MM level of theory. A spherical droplet was extracted to compute two-dimensional B3LYP/6-31G**/MM potential energy surfaces (PES) in both forward and reverse directions. Selected reaction coordinates for the phosphoryl transfer are shown in Fig. 2 B and those for the crosslink of Ser861 with Remdesivir are shown Fig. 3 B, (see also Supplementary Fig. 4 and Supplementary Methods). The AMBER program interfaced with Terachem 1.9^25, 26^ was used for the QM/MM calculations. Electrostatic embedding was used in all hybrid calculations.

**Fig. 4.**
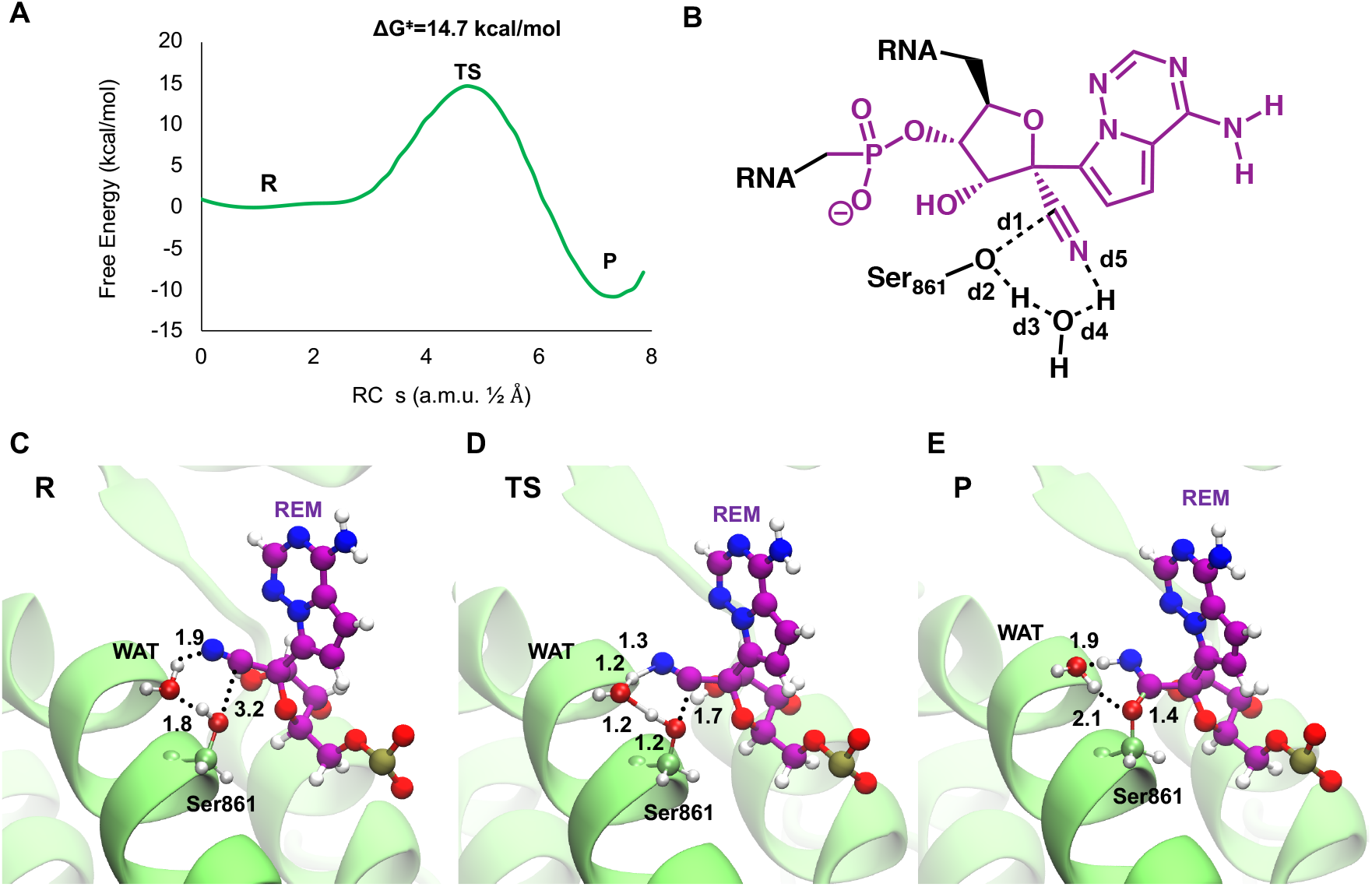
Remdesivir forms a covalent complex with RdRp of SARS-CoV-2. **A** Free energy profile for the attack of Ser_861_ to the C atom of nitrile group of Remdesivir (see the bidimensional PES in Supplementary Fig. 11). **B** Scheme representing the concerted TS where nucleophilic attack of Ser_861_ occurs at the same time that a water mediated proton transfer to the nitrile group of Remdesivir. Distances included in the definition of the Reaction Coordinate (s) are shown. **C** Active site insight of Reactant state. Distances involved in the reaction are shown as dotted lines with its average value in Å. **D** Active site insight of the the TS. **E** Active site insight of the covalent complex formed by Remdesivir and RdRp.

### QM/MM Exploration of the Minimum Free Energy Paths and Potential of Mean Force

QM/MM-MD simulations were performed to obtain minimum free energy paths (MFEPs) by means of the string method.^27, 28^ This method allowed us to explore different reaction mechanisms and select the preferred one in terms of free energy. From 60 to 120 string nodes were used. Afterwards, a path collective variable (CV)^28^ was defined to obtain the potential of mean force (PMF) using from 60 to 120 umbrella sampling^29^ windows. The chosen set of CVs that followed the progress of the reactions, the breaking and forming bonds, are shown in Fig. 2 B, Fig. 3 B and Supplementary Methods (see Supplementary Fig. 4). MFEPs were obtained at the DFTB3^30, 31^/MM level and the PMFs were corrected at the B3LYP/6-311++G** level. The error of the free energy barriers was calculated as 95% confidence intervals and reached values within ± 1 kcal·mol^−1^. Each of the sampling windows consisted on 20ps of equilibration followed by 200ps of production. The AMBER program^15^ with electrostatic embedding was used for the QM/MM calculations. Corrections at the high level of theory were made with Gaussian16^32^ (see Supplementary Methods).

### MD simulations on nascent RNA

We performed MD simulations on two duplexes: r(CGCGAAUUGCGC)·r(GCGCAAUUCGCG) and r(CGCGARUUGCGC)·r(GCGCAAUUCGCG) to determine the structural impact of the introduction of a Remdesivir in a canonical RNA duplex. Starting structures were those expected for a canonical RNA duplex as implemented in AMBER. Systems were hydrated, minimized, thermalized and equilibrated using standard protocols^33, 34^ before 1 μs long MD trajectories at constant temperature (T=298K) and pressure (P=1 atm). Details of simulations are shown in Supplementary Methods.

### Sequence analysis

Mutations affecting RdRp were reconstructed using the augur pipeline to infer nucleotide changes at the internal nodes^35^ applied to 12397 GISAID samples obtained on 30/05/2020. The consequence type of the RdRp mutations was annotated using a customized implementation of the Ensembl Variant Effect Predictor (VEP version 92) using the first SARS-CoV-2 sequenced genome (NCBI ID: NC_045512v2) as a reference. Afterwards, mutations were mapped on the MD-refined structure of the RdRp-RNA complex to locate their proximity to crucial functional regions. The aggressiveness of each individual mutation was evaluated from BLOSUM80 matrices.^36^ Homolog search were performed with BLASTP^8^ as implemented in the ncbi web server (https://blast.ncbi.nlm.nih.gov/). Alignments were performed using EMBOSS program^37^ as implemented in the EBI webserver (https://www.ebi.ac.uk/Tools/psa/).

## RESULTS AND DISCUSSION

### SARS-CoV-2 RdRp active site suggests good fidelity and activity

The viral protein (see Fig. 1 A) has a quite canonical active site similar to those of other polymerases. Two essential Mg^2+^ coordinate the β and γ phosphate groups of the incoming triphosphate nucleotide as well as Asp_618_, Asp_760_, Tyr_609_, and the O3’ terminal of the negative RNA strand (see Fig. 1 B), which according to the circular reaction mechanism for polymerases by de Vivo and coworkers is expected to be ionized.^38^ Two Arginine residues (Arg_624_ and Arg_553_) coordinate the phosphates of the incoming nucleotide aligning the gamma phosphate for an effective transfer (see Fig. 1 C). The affinity for ribonucleotides triphosphate substrates can be explained by the need for North puckering (otherwise reactive groups are not well aligned), as well as by a myriad of specific H-bonds between the 2’OH group of the NTP and the catalytic site Asp_623_ and Ser_683_ side chains (see Fig. 1 C). Additional hydrogen bonds are found between i+1 2’OH and Ser_759_ (see Fig. 1 D). The base-specificity is controlled by the complementarity of hydrogen bonding with the template nucleobase (see Fig. 1 E, Supplementary Fig. 5), by the phosphate coordination, as well as by the residues surrounding the active site that embrace the pairs introducing strong isosteric requirements that make very unlikely a non-Watson-Crick pairing scheme (see Fig. 1 E). Overall the active site emerging from EM structures^12,14,39,40^ and atomistic simulation strongly suggest that despite the reduced evolutionary refinement, SARS-CoV-2 RdRp has all the structural requirements to be an efficient RNA polymerase, both in terms of catalysis and substrate specificity.

### The RNA-pol reaction mechanism

QM/MM simulations were used (see Methods and Supplementary Methods) to study the ability of SARS-CoV-2 RdRp to incorporate a natural tri-phosphate (NTP, exemplified here by UTP), as well as RTP (Remdesivir-TP) into a nascent RNA. Careful analysis of trajectories and of the free energy profiles shows that the calculation is well converged leading to a smooth free energy profile with a single maximum (Transition State, TS), a free energy of activation of 15.8 kcal·mol^−1^ for UTP (see Fig. 2, Supplementary Fig. 6 and Supplementary Video 1) and a negative free energy of reaction (around 5 kcal/mol prior to PPi release). Very interestingly, kinetic parameters for RdRp compare well to those associated to prokaryotic or eukaryotic polymerases,^41–43^ which combined with the similarity in the reaction mechanism allow us to conclude that the enzyme can be quite novel in evolutionary terms, but it is as efficient as the highly evolved eukaryotic polymerases.

Trajectories show that RTP fits very well into the active site showing strong canonical Watson Crick interactions with the uridine in the template RNA. Its isosterism with adenosine allows a perfect shape complementarity at the binding site showing an arrangement of reactive groups that predicts that it will be a substrate rather than an inhibitor. This hypothesis is confirmed by accurate QM/MM simulation that show that incorporation of RTP can happen with a free energy barrier only slightly larger than that of a natural substrate (10% up to 17.4 kcal·mol^−1^; see Fig. 2, Supplementary Fig. 7 and Supplementary Video 2). Such an increase is due mostly to a slight misalignment of the Oα-Pα-O3’ attack angle (161 ± 8° for RTP and 172 ± 5° for UTP; see also Supplementary Fig. 6-8). Thus, in agreement with recent EM experiments14 we can rule out the possibility of Remdesivir inhibiting RdRp by blocking the ATP-binding site of RNA polymerase. On the contrary, our simulation strongly suggests that RTP can be efficiently incorporated in front of uridine in a nascent RNA strand.

### Remdesivir is well tolerated in an RNA duplex

Remdesivir is a C-nucleoside that may produce distortions in the helix which might result in a delayed inhibition of the enzyme due to incorrect displacement of the nascent duplex along the exit channel. To explore this possibility, we performed MD simulations of two RNA duplexes differing only in the substitution of a central r(A·U) pair by r(R·U) one (see Methods). Results summarized in Fig. 3 strongly suggest that Remdesivir is well tolerated in an RNA duplex and does not introduce any major structural distortion which could justify termination of RNA synthesis. Particularly, there are not significant differences between the hydrogen bonding stability of r(A·U) and r(R·U) pairs (see Fig. 3 C). With the required cautions derived from the well-known uncertainties of RNA force-field and of the limited sampling (1 μs), our results argue against the idea that a dramatic structural alteration of the RNA duplex is the key responsible for Remdesivir-induced termination of the RNA synthesis.

### Remdesivir can crosslink RNA and RdRp

Trying to explore alternative reasons for the inhibitory properties of Remdesivir we slide the nascent RNA duplex along the exit tunnel of RdRp to make the r(R·U) pair by simulating addition of extra nucleotides. This allows us to scan interactions of the RNA at several positions along the exit tunnel. After R was incorporated we were not able to detect any point of steric clash that could justify stopping the polymerase progression (see Supplementary Fig. 9). Interestingly, when three more nucleotides were incorporated, we found a Ser_861_ whose side chain locates at around 3.7±0.3 Å from the nitrile group of Remdesivir (see Supplementary Fig. 10 A B). The distance does not justify however a steric clash considering the large flexibility of the Ser sidechain, especially in a well-solvated microenvironment (see Supplementary Fig. 10 A), arguing against the hypothesis that nitrile-Ser_861_ steric clash can explain the inhibitory properties of Remdesivir.^10^ However, a close look to the trajectories shows that the Ser_861_ side chain is frequently pointing towards the nitrile group of Remdesivir in an arrangement suitable for a nucleophilic attack and in the presence of water that spontaneously enters into the cavity (see Supplementary Fig. 10 C D). In summary, it is placed in an optimal position as to act as a general acid/base catalyst. Note that, while not suggested for this protein, attacks of serine and cysteine to nitrile groups have been characterized for other proteins,^44, 45^ which drives us to explore in more detail this potential reaction mechanism by using a QM/MM strategy similar to that used to describe the polymerization reaction (see Methods). Results (see Fig. 4, QM/MM PES in Supplementary Fig. 11, and Supplementary Videos 3 and 4) show a concerted reaction mechanism where, while hydroxylic group of Ser_861_ attacks the carbon atom of the nitrile group, a water molecule (see Supplementary Fig. 10 D) abstracts one proton from the hydroxyl group of Ser_861_, donating it to the N atom of nitrile group (see Fig. 4 D, Supplementary Video 3). The structures sampled along the minimum free energy profile agree with the electronic flow for a Pinner reaction.^46^ Our calculations suggest an exergonic (−10.7 kcal·mol^−1^; see Fig. 4 A) process showing a single TS (as expected from a concerted mechanism; see Figure 4 and Supplementary Fig. 11) with an activation barrier of 14.7 kcal·mol^−1^; see Fig. 3 A), which compare with the activation barrier of polymerization of a UTP (15.8 kcal·mol^−1^) and RTP (17.4 kcal·mol^−1^).

QM/MM calculations suggest the existence of a kinetic competition between UTP incorporation and crosslinking leading to enzyme inhibition. In principle, barrier for UTP incorporation is slightly larger than that of crosslinking, which suggests that the RNA will be kinetically trapped stopping the extension of the RNA, and inhibiting then the activity of the enzyme. However, we can expect residual polymerization leading to longer transcripts, as experimentally observed.^9^ A larger concentration of Remdesivir might favor the inhibition by a double mechanism: by increasing the probability of RTP incorporation into nascent RNA and also by enlarging the gap in relative activation free energy between polymerization and crosslinking (from 1.1 kcal·mol^−1^ to 2.7 kcal·mol^−1^), and driving then the kinetic balance towards the inhibition side.

The proposed covalent inhibition mechanism is consistent with the incorporation of three more nucleotides after incorporation of Remdesivir to RNA^9^ and agrees with a myriad of indirect evidences other than the delay inhibitory properties of Remdesivir. For example, mutational analysis shows that Ser_861_ has a crucial role in R-induced inhibition of RdRp.^40^ Furthermore, large and systematic efforts to substitute the nitrile group by less problematic substituents (the nitrile group appears as a “probably undesired” consequence of the Remdesivir synthetic pathway) resulted in molecules with a lower inhibitory profile.^47^ Finally, Pinner reaction has not been described for RdRp, but has been experimentally validated for many several nitrile-containing drugs designed to inhibit cysteine or serine proteases.^48, 49^ Altogether, results strongly suggest that Remdesivir inhibits SARS-CoV-2 by a covalent mechanism.

### Mutational analysis of SARS-CoV-2 RNA-pol agrees with the proposed reaction mechanisms

Thousands of SARS-CoV-2 viruses have been sequenced, showing that RdRp accumulates a non-negligible number of mutations, which lead to viable virus, i.e., to active RdRp. We should then check whether some of these mutations invalidate our reaction mechanism. As shown in Supplementary Fig. 12, this is not the case, missense variant leading to amino-acid changes are typically mild according to BLOSUM80 matrices and are located in loops on the surface of the protein (Supplementary Fig. 12 A). Particularly, the active site and the exit tunnel are very well conserved during the viral evolution (see Supplementary Fig. 12 B, C). Moreover, virus has not explored yet mutational landscape in the region of the Ser_861_ (see Supplementary Fig. 12 D).

### Remdesivir is not likely to affect human RNA polymerases

Remdesivir enters in human cells and is transformed into RTP also by human enzymes. Considering the similarity in the mechanism of reaction of SARS-CoV-2 RdRp and human polymerases we could expect Remdesivir being incorporated into RNA, which could rise to inhibition of the human enzyme with toxic consequences. To check this possibility we explored the extension of the nascent RNA along the exit channel to trace close interactions between the nitrile group and serine or cysteine residues in the two crystallized human RNA polymerases (RNA polymerase^50^ and human RNA polymerase II^51^). We did not find any of these residues in the expected displacement path of Remdesivir once incorporated into the nascent RNA (see Supplementary Fig. 13). In other words, no significant inhibition of the human polymerases is expected. Note that this explains the reduced toxicity of Remdesivir in humans.^52, 53^

### Inhibition mechanism can be common to other viruses

Ser_861_ is highly conserved in other CoVs RdRps (see Supplementary Fig. 14), placing a constant position in an alpha helix that adopts a highly conserved 3D arrangement (compare SARS-CoV-2 and SARS-CoV structures)^12,14,39,40,54^ and this position is well preserved in those cases for which CoV RdRps structure is available. This would suggest that Remdesivir might be effective against many other CoV RdRps. The mechanism of inhibition of other distant RNA-viruses (Marburg virus, Ebola, Hepatitis C and many others) might be similar, but needs to be elucidated. It is tempting to believe that enzymes evolved to discriminate between DNA (2’deoxyribose) and RNA (ribose) might be decorated with a large number of serine, tyrosine or threonine in the exit channel, suggesting a general delay inhibitory mechanism like that suggested here. Note that this would agree with the delayed inhibition of Ebola RdRp by Remdesivir.^55^ The lack of structural information and the poor sequence similarity between the RdRps precludes to confirm this hypothesis.

## CONCLUSIONS

SARS-CoV-2 RdRp is a protein common to other coronaviruses, but showing little homology out of the family. As CoVs are just a few thousand years old, the protein has had a limited evolutionary period and we could expect low efficiency. However, our calculations show that it has a well refined reaction site able to select the entering nucleotide and to catalyze its addition to a nascent RNA. The viral enzyme follows a mechanism that is similar to that of bacterial or eukaryotic polymerases with the transferred phosphate being stabilized by 2 Mg^2+^ ions exquisitely coordinated by acidic residues of the catalytic site, while the phosphates of the incoming nucleotide being stabilized by a network of basic residues.

Our simulations very clearly suggest that RTP is not inhibiting the active site by competing with ATP at the catalytic site, but that it is incorporated into the nascent RNA in front of a uridine. The resulting duplex does not show dramatic structural changes which would hinder displacement of the nascent duplex along the exit channel. In fact, analysis of the displacement of the RNA-containing Remdesivir along the exit tunnel fail to detect points of steric clashes, but show the spontaneous formation of a catalytical arrangement that justifies a Pinner reaction (between the nitrile group of Remdesivir and Ser_681_), leading to the formation of a protein-RNA covalent bond. However, we cannot determine whether this inhibition will be reversible or not, as inspection of the free energy curve (Fig. 4D) and the behavior of similar inhibitors^49^ for other proteins suggest that it may display slow reversibility.^49, 56^ This potential reversibility combined with the kinetic competition between crosslinking and polymerization might explain the unusual inhibitory properties of Remdesivir. Results presented here open then the possibility to design better inhibitors of CoVs RdRps and to a rethinking on the use of covalent inhibitors of pathological proteins.

## Acknowledgments

This work has been supported by the Spanish Ministry of Science (BFU2014-61670-EXP), the Catalan SGR, the Instituto Nacional de Bioinformática, and the European Research Council (ERC SimDNA), the European Union’s Horizon 2020 research and innovation program under grant agreement No 676556, the Biomolecular and Bioinformatics Resources Platform (ISCIII PT 13/0001/0030) cofunded by the Fondo Europeo de Desarrollo Regional (FEDER), and the MINECO Severo Ochoa Award of Excellence (Government of Spain) (awarded to IRB Barcelona). M. O. is an ICREA academia researcher. J. A. acknowledges the Spanish Ministry of Science for a Juan de la Cierva post-doctoral grant. J. A. thanks Dr. K. Zinovjev for the string-method technical support. J. A. thanks the support team of MareNostrum supercomputer and support team at IRB Barcelona. Calculations were performed on the MareNostrum supercomputer at the Barcelona Supercomputer Center and in MMB cluster. M. O. and J. A. thank Prof. Núria López-Bigas and Dr. Francisco Martínez-Jiménez for their help in filtering data of mutations.

