## Supplementary Information for "RNA-Dependent RNA Polymerase From SARS-CoV-2. Mechanism Of Reaction And Inhibition By Remdesivir"

J. Aranda[1], M. Orozco[1,2]*

[1] Institute for Research in Biomedicine (IRB Barcelona), The Barcelona Institute of Science and Technology, Baldiri Reixac 10, 08028 Barcelona, Spain
[2] Departament de Bioquímica i Biomedicine, Universitat de Barcelona, Universitat de Barcelona, Avinguda Diagonal 645, 08028 Barcelona, Spain

* Correspondence to
M.Orozco.

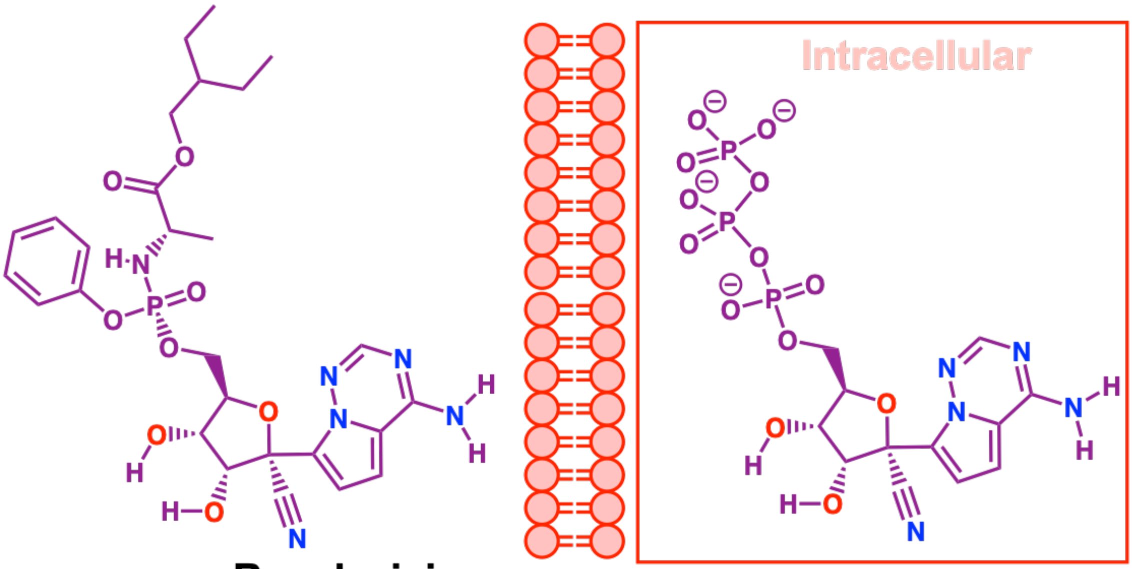


Remdesivir

Remedesivir-TP

**Supplementary Fig. 1.** Chemical structures of Remdesirvir in its prodrug form and as a triphosphate nucleoside inside the cell.

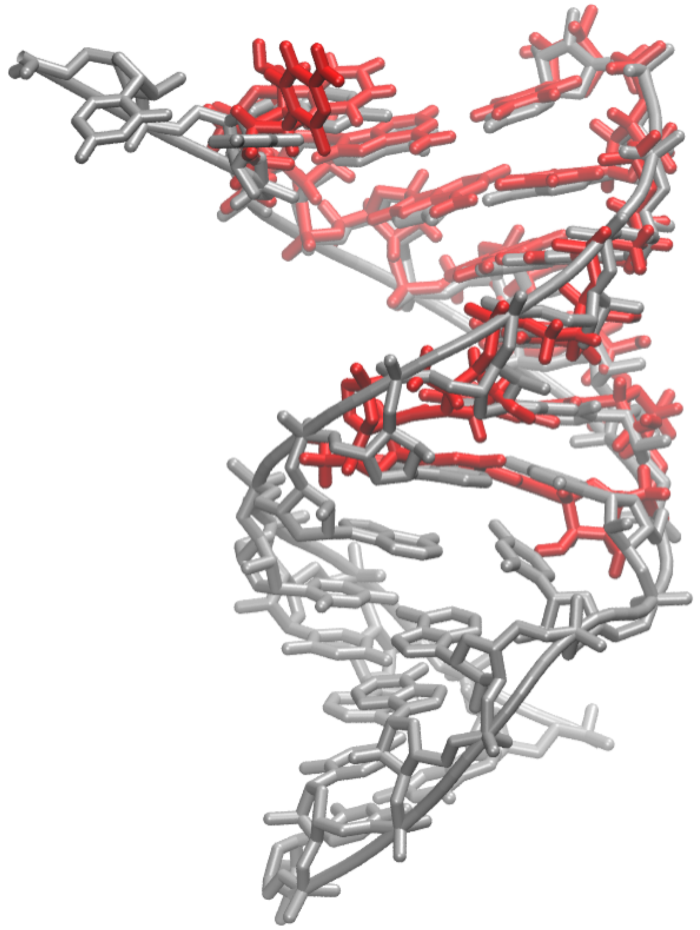

**Supplementary Fig. 2.** Alignment of RNA from cryo-EM structure with PDB ID:7bv2,[1] displayed in gray, and the double stranded RNA optimized and employed in our simulations and extracted from Hepatitis C virus, PDB ID: 4wtg,[2] displayed in red. RMSD between backbone atoms of the nucleic acids is 1.1 Å.

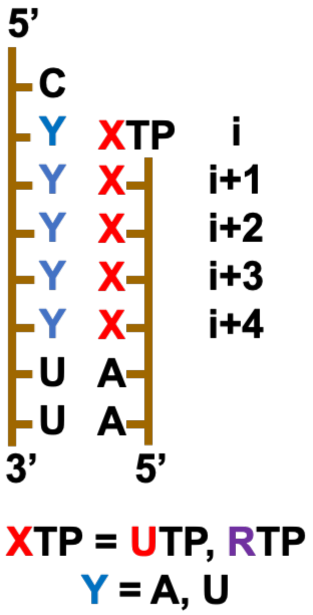


**Supplementary Fig. 3.** RNA templates used in this study. One template consisted on natural occurring nucleosides, the rest contained only one Remdesivir in the depicted positions of the nascent strand. In this way we moved the r(R·U) pair from i to i+4. X refers to a U or a Remdesivir (R) which are paired with A or U respectively.

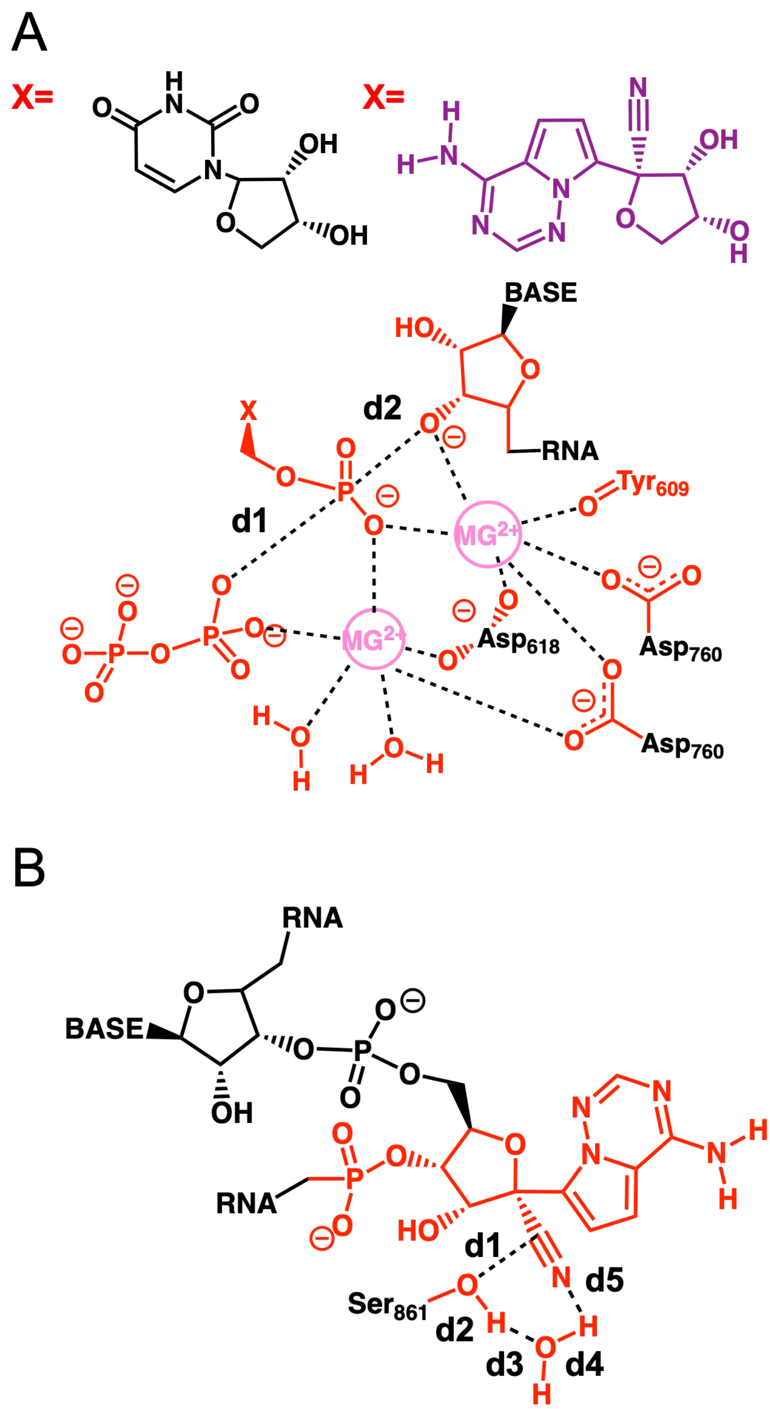


**Supplementary Fig. 4 A** Atoms described at QM level (in red and pink) in the hybrid QM/MM calculations during the ligation reaction step. Distances involved in the Reaction Coordinates employed are shown. **B** Atoms described at QM level (in red) in the hybrid QM/MM calculations for the step where a $Ser_{861}$ nucleophilically attacks the nitrile group of Remdesivir. Distances involved in the Reaction Coordinates employed are shown.

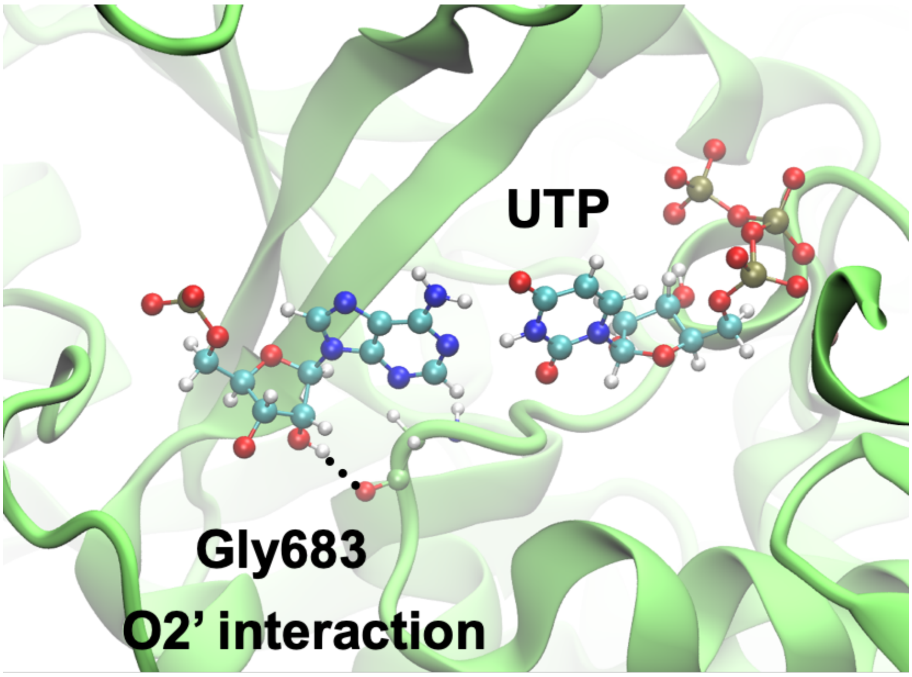


**Supplementary Fig. 5.** $Gly_{683}$ forms a hydrogen bond interaction with the O2' hydroxyl group of the template, recognizing the entry of a ribonucleotide template.

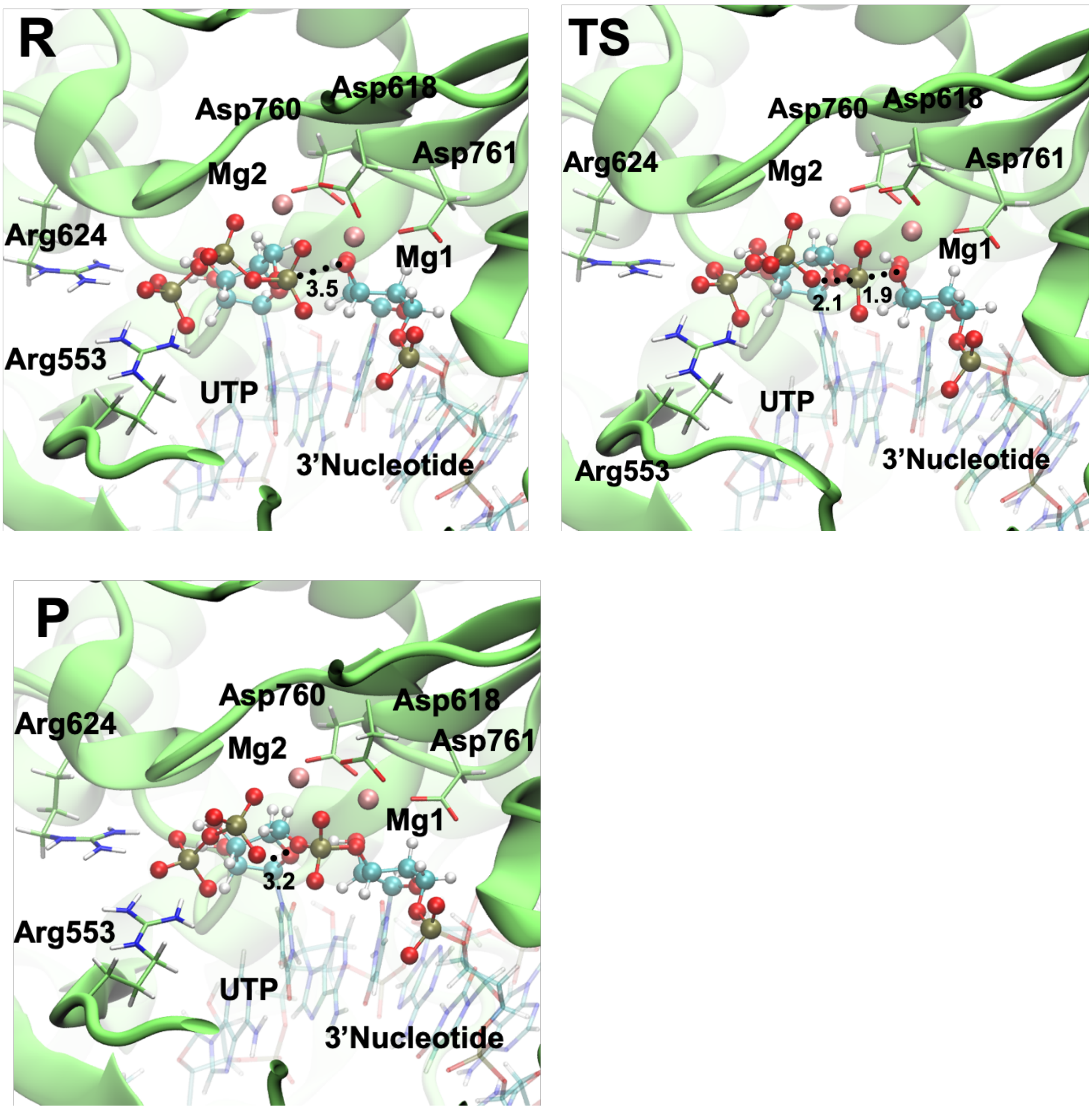


**Supplementary Fig. 6.** Active site insights of RdRp when a U is being incorporated to a nascent viral RNA strand. Reactant, Transition and Product states representative structures are depicted. Distances involved in the phosphoryl transfer are shown in Å and depicted as dotted lines.

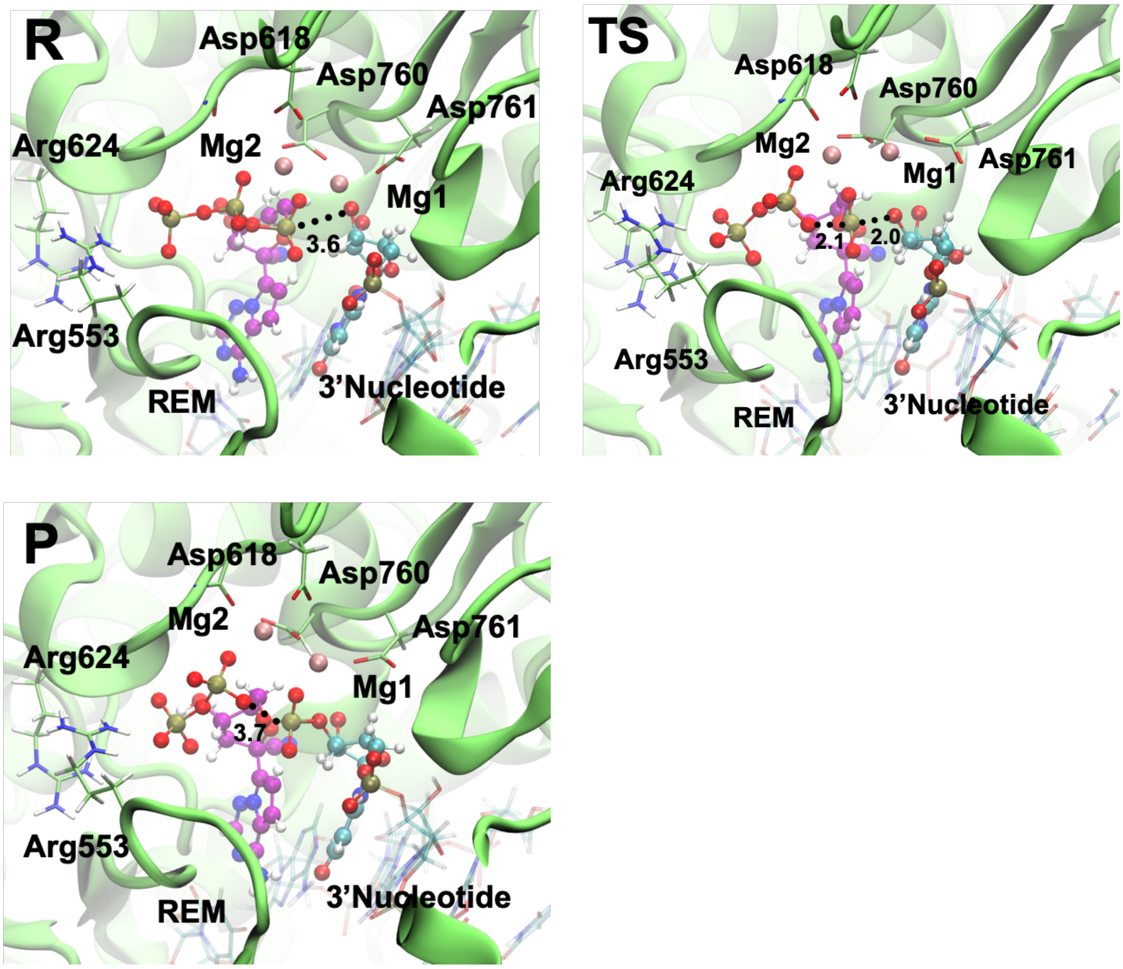


**Supplementary Fig. 7.** Active site insights of RdRp when a Remdesivir is being incorporated to a nascent viral RNA strand. Reactant, Transition and Product states representative structures are depicted. Distances involved in the phosphoryl transfer are shown in Å and depicted as dotted lines.

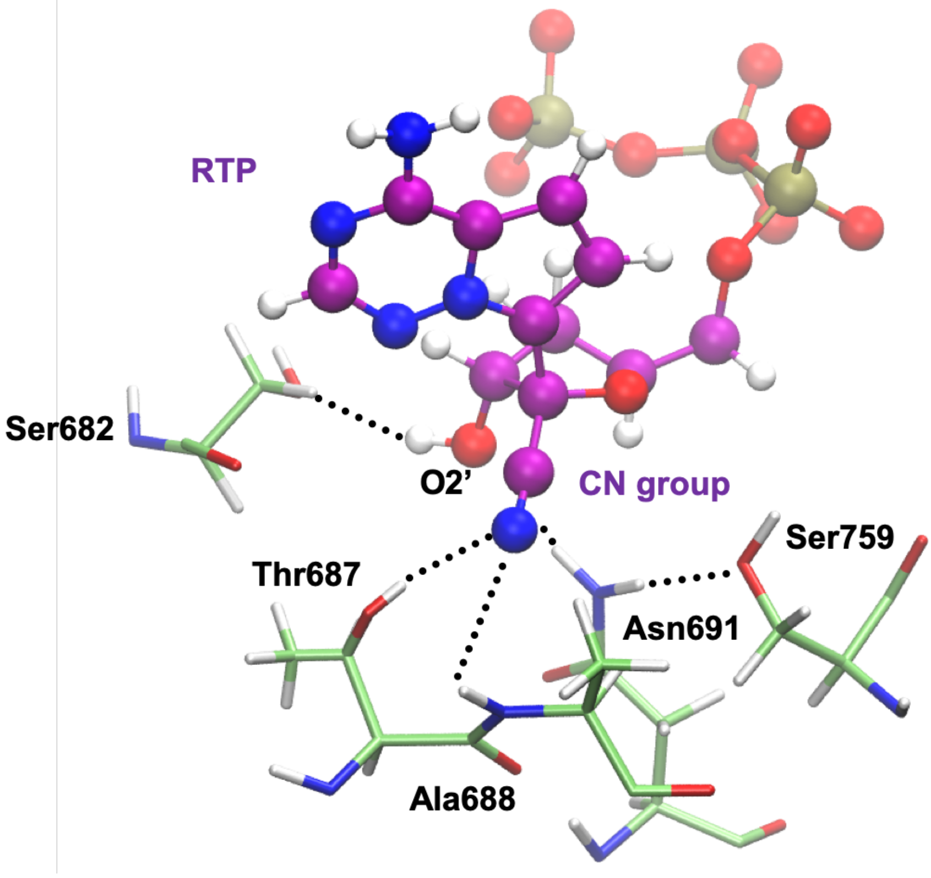


**Supplementary Fig. 8.** Interactions that are formed between the nitrile group of Remdesivir-TP (RTP) and the active site of RdRp of SARS-CoV-2 during our simulations. Remdesivir C atoms are shown in purple. Most important interactions are depicted as dotted lines. These interactions place Remdesivir nitrile group towards the cavity and in a slightly different arrangement than UTP. Same pattern of recognition of the O2’ group of RTP is achieved through the $Ser_{682}$ residue.

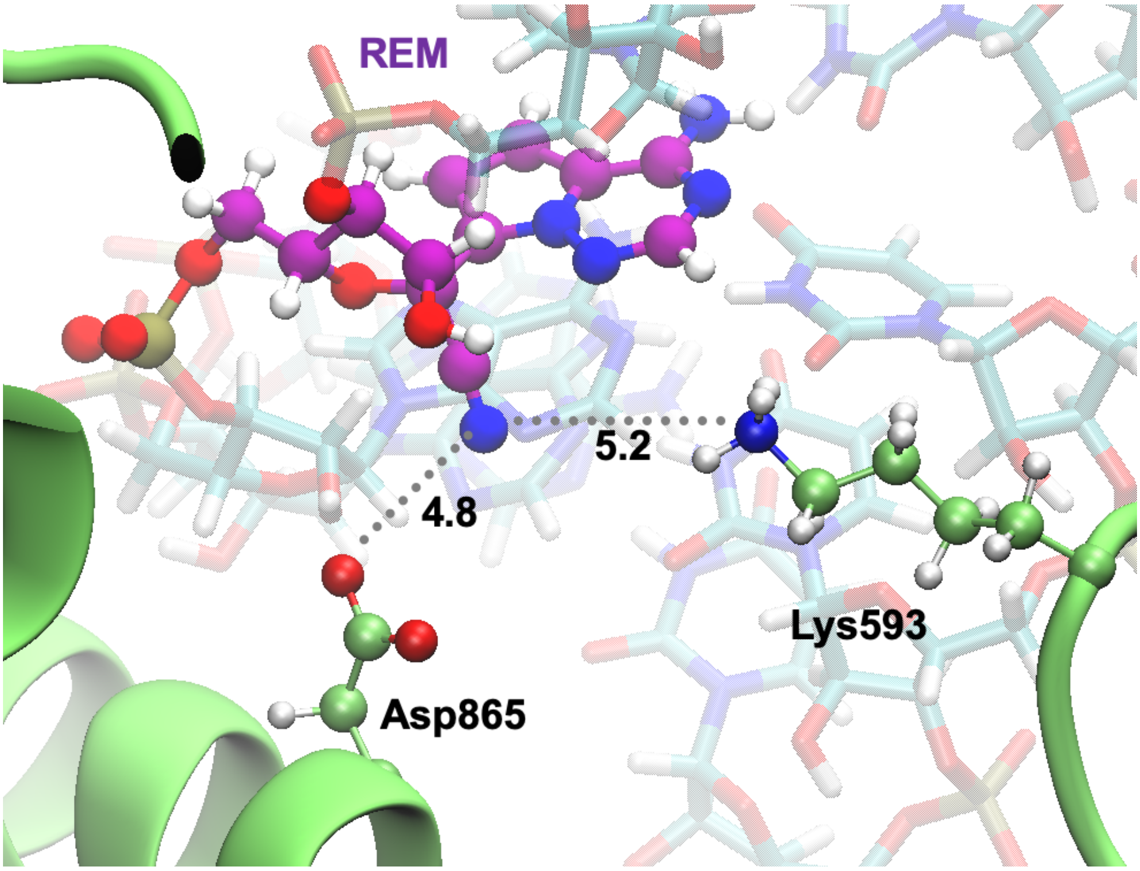


**Supplementary Fig. 9.** Closest residues of RdRp to Remdesivir after two nucleotides have been incorporated. Residues are far to interact with any polar atom of Remdesivir. Average distances during MD simulation between $Asp_{865}$ and $Lys_{593}$ sidechains to N atom of nitrile group of Remdesivir are shown in Å.

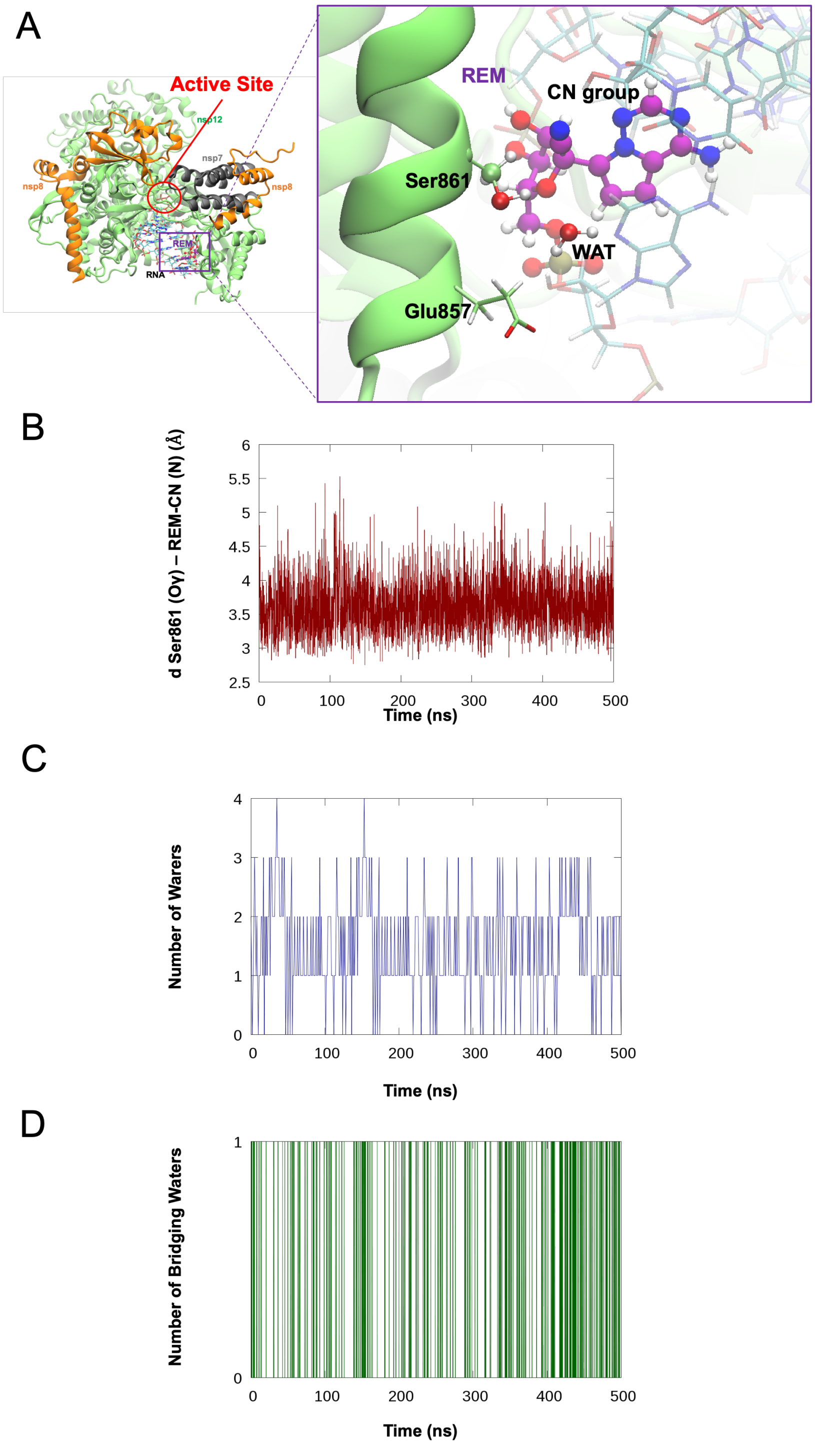


**Supplementary Fig. 10. A** Insight of the surroundings when Remdesivir CN group is close to Ser$_{861}$. **B** Ser$_{861}$ is found close to the nitrile group along 500 ns of MD simulation. **C** Number of waters within 3.5 Å distance from nitrile group. **D** Number of waters hydrogen bonded at the same time with the N atom of nitrile group of Remdesivir and the sidechain Oγ atom of Ser$_{861}$.

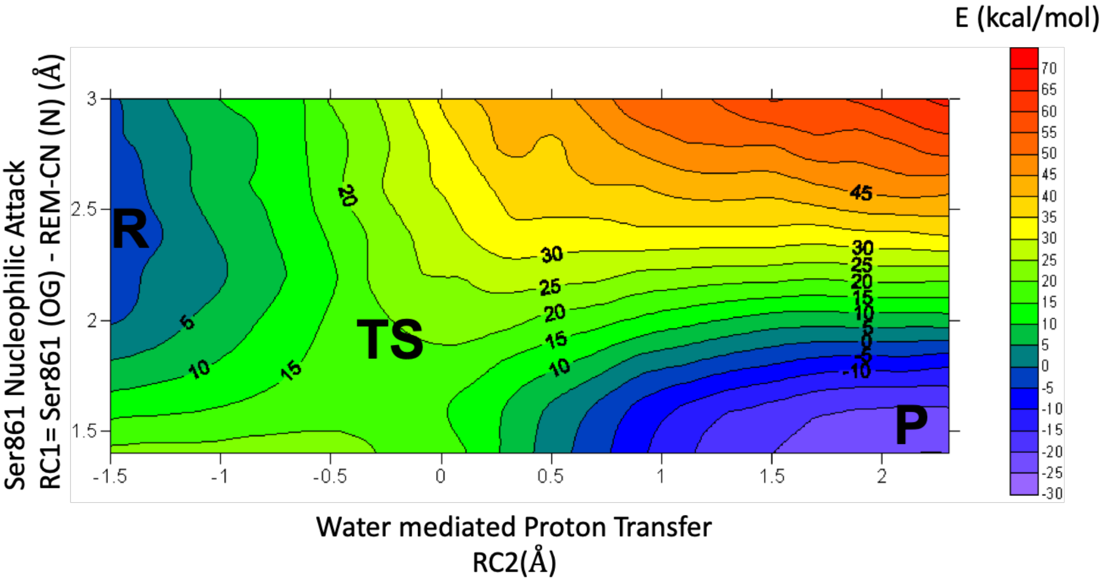


**Supplementary Fig. 11.** Potential Energy Surface calculated at the B3lyp/6-31G**/MM level monitoring the possibility of a nucleophilic attack of $Ser_{861}$ to C atom of nitrile group of Remdesivir (Reaction Coordinate 1, RC1=d1 as shown is Supplementary Fig. 4 B), and a water mediated proton transfer from $Ser_{861}$ to a water molecule which in turn donates a proton to the N atom of nitrile group of Remdesivir (RC2=(d2-d3) + (d4-d5)), as shown is Supplementary Fig. 4 B). To further details we refer the reader to the supplementary methods section. A movie depicting the minimum energy path along the PES from reactants to products in provided (see Supplementary Video 4).

A

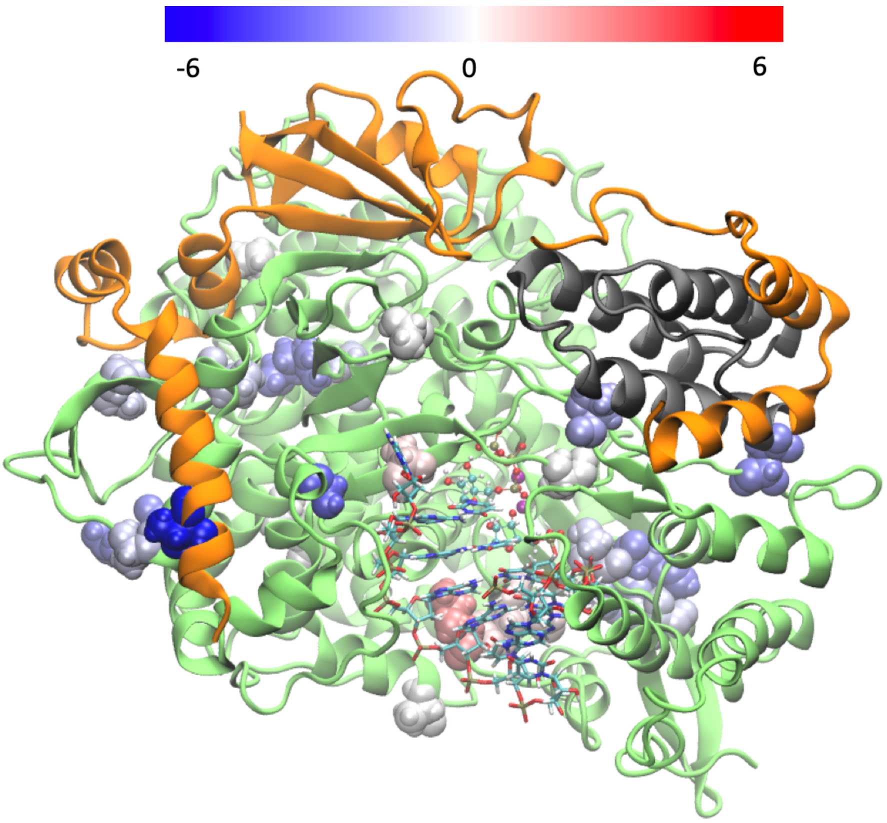

-6
0
6


B

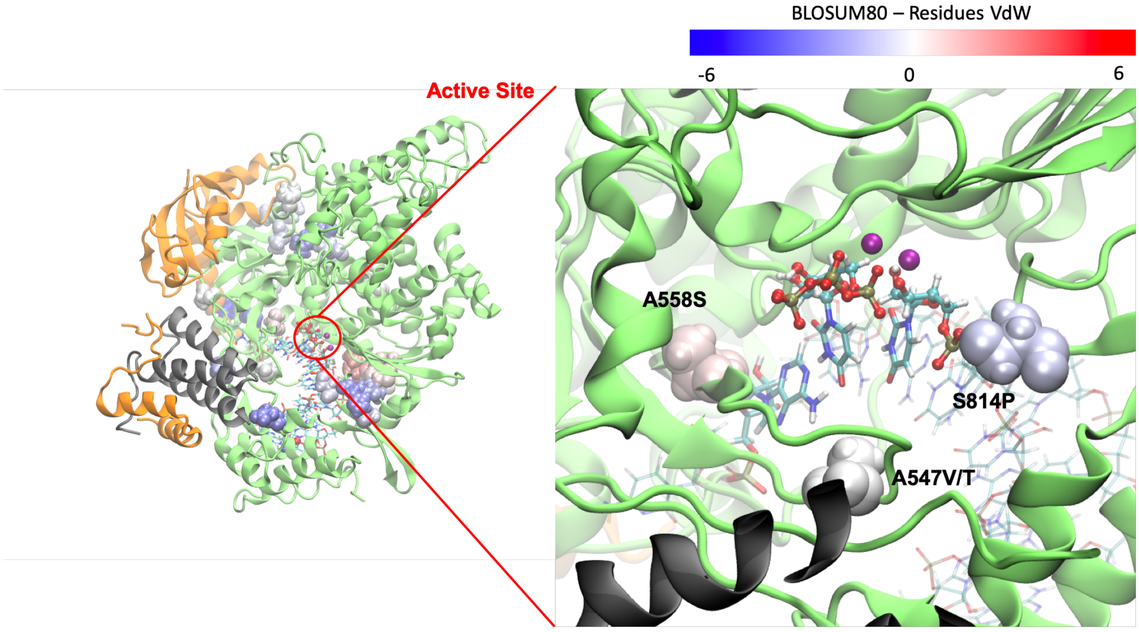

BLOSUM80 – Residues VdW
-6
0
6
Active Site
A558S
S814P
A547V/T

C

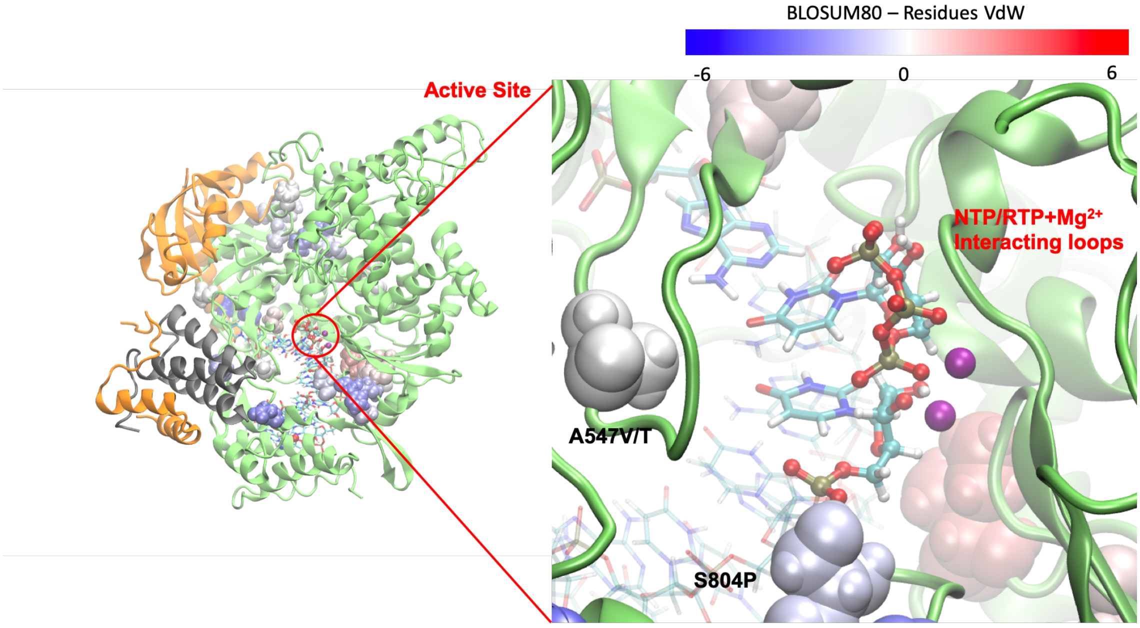


D

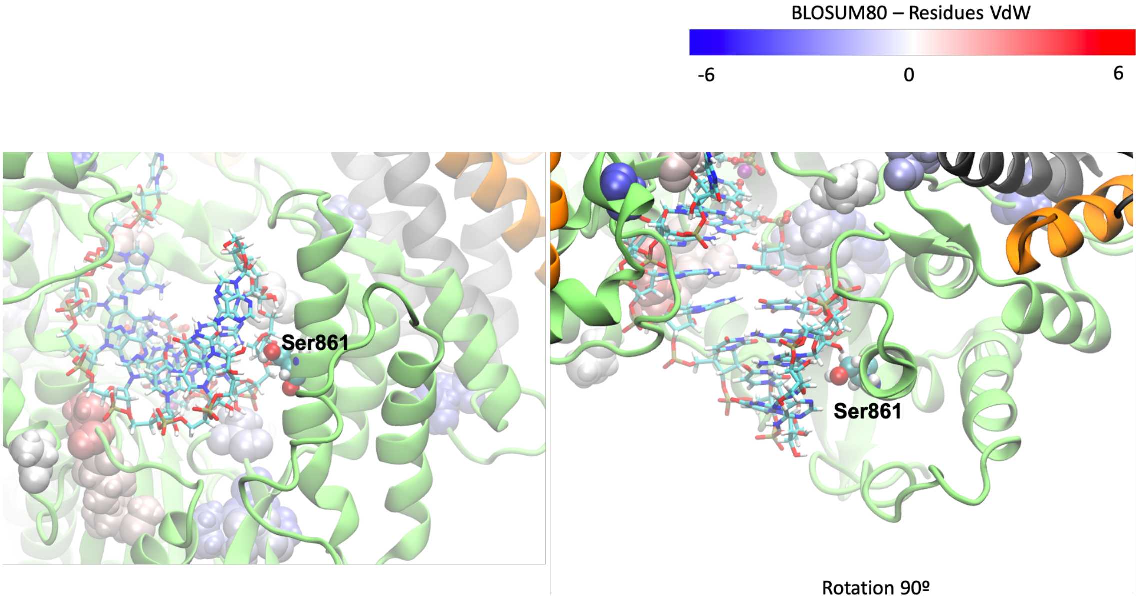


**Supplementary Fig. 12 Mutational analysis of RdRp of SARS-CoV-2. A** Overall view of RdRp-RNA complex. Most important residues which suffer from mutation are depicted. Residues are in VdW representation and color based on its BLOSUM80 scoring (blue or -6 is the most disruptive score found, and red or 6 was the most favorable). **B** Mutations close to the active site of RdRp. **C** No mutations are found in the interacting loops with NTP and $Mg^{2+}$ ions. **D** No mutations are found in the vicinities where $Ser_{861}$ is located.

A

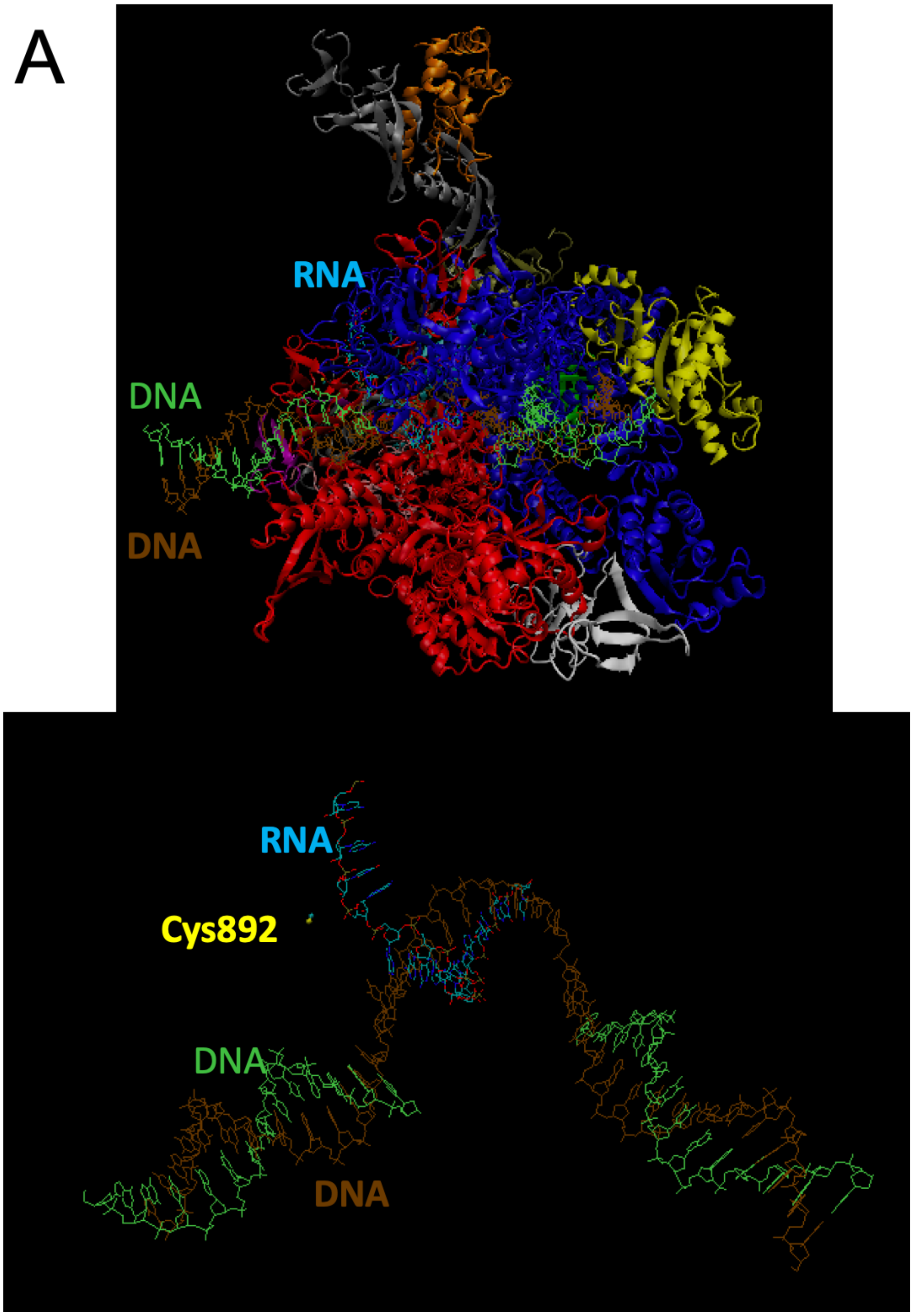

B

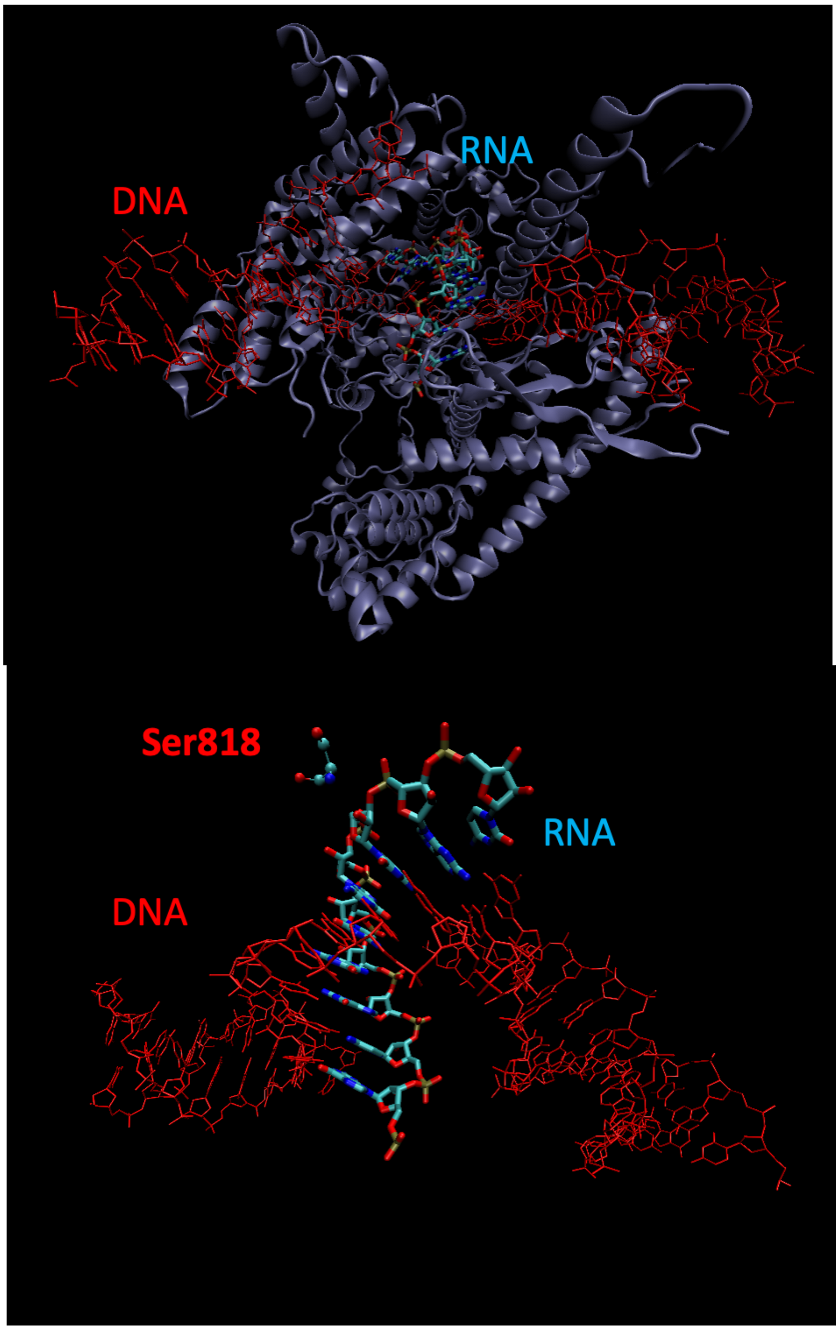


**Supplementary Fig. 13 A** Human RNA polymerase II-DNA/RNA complex (PDB ID: 5FLM[3]). $Cys_{861}$ which is the closest Cys or Ser residue to a C1’ atom of the nascent RNA strand, is placed to more than 8 Å distance. **B** Human mitochondrial RNA polymerase in complex with DNA and RNA, PDB ID: 4BOC.[4] $Ser_{818}$, the closest Cys or Ser residue to a C1’ atom of the nascent RNA strand is placed at 6.3 Å distance.

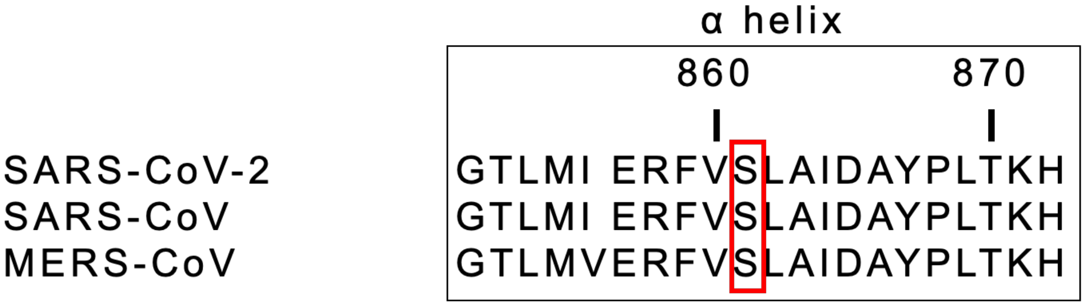


**Supplementary Fig. 14** Alignment of SARS-CoV-2, SARS-CoV and MERS-CoV sequence for the alpha helix where $Ser_{861}$ is placed.

### SUPPLEMENTARY METHODS

#### *System set up*

Our starting point was the recently published cryo-EM structure of the of the SARS-CoV-2 RdRp in complex with its cofactors.[5] As this structure was resolved without the $Mg^{2+}$ cations needed for the catalysis and without the RNA template and nascent strand, we aligned it with the Hepatitis-C virus X-ray structure[2] which was crystallized with two $Mn^{2+}$ cations, a nucleotide analog diphosphate molecule and a RNA template strand and a nascent RNA strand. We selected this structure as it showed the best alignment for both the cleft where RNA binds and the active site. In addition, it was resolved with two catalytic divalent cations and a diphosphate nucleotide analogue, which enabled us to build our selected substrates based on X-ray positions. Alignment was performed making use of the Pymol program, selecting a set of atoms that consisted on the atoms in the catalytic domains, and the residues placed in the cleft which are in charge of the RNA binding of both RdRp's. Thus, we then used the RNA molecule as well as the two cations and the diphosphate nucleotide molecule in our SARS-CoV-2 RdRp systems. The two cations were modeled as $Mg^{2+}$ cations, and the nucleotide diphosphate was used as a template to build the UTP or Remdesivir-TP molecules. Afterwards the systems were protonated making use of the LEAP module of the AMBER program.[6] The systems were solvated with LEAP module into a truncated octahedron box of TIP3P water molecules with a buffer of water molecules extending for 12 Å in every direction around the systems. Systems were neutralized by adding K+ ions. For the magnesium ions the parameters developed by Allner et al. were employed.[7] Proteins were described with ff14SB[8] AMBER ff. The RNA was simulated by combining ff99, the PARMBSC0 modifications and the chiOL3 modifications for RNA.[9–12] Charges and parameters for the non-standard residues were derived to be compatible with the employed AMBER force field making use of the RED server.[13] Specifically, we derived parameters for a 3'-terminal uridine nucleotide deprotonated at its O3', a 3'-terminal Remdesivir nucleotide deprotonated at its O3', a Uridine-TP, a Remdesivir-TP and a Remdesivir nucleotide.

***Molecular Dynamic Simulations***

Classical MD simulations were carried out using the AMBER 18 program[6] with a time step of 2 fs and applying the SHAKE algorithm[14] to bond lengths involving hydrogen atoms. Simulations were carried out in the isothermal-isobaric ensemble with a pressure of 1 atm and a temperature of 298 K. The Berendsen algorithm[15] was applied to control the pressure and the temperature with a coupling constant of 5 ps. The Particle Mesh Ewald method[16] was used to compute long-range electrostatic interactions using standard defaults and a cutoff in the real-space of 10 Å. The systems were energy minimized, thermalized and pre-equilibrated for 100 ns before the production run was conducted. During this multi-step approach, we firstly equilibrated the water box and counterions, then released the side-chains of the protein residues and then released the nucleobases gradually by maintaining its backbone frozen. Afterwards we released the whole protein atoms by maintaining the active site residues (UTP or RTP, the $Mg^{2+}$ cations and their coordination spheres) and the nucleic acid backbone frozen. We then performed an MD run imposing a restraint to the distance between the 3'-hydroxyl oxygen atom of the terminal nucleotide and the α-phosphate atom of UTP or RTP. Finally, we slowly released the positional restraints imposed to the system and the distance restraint. A total time of 500 ns of fully unrestrained MD simulations were performed for all the systems. Thus, we performed MD simulation in systems containing RdRp with its cofactors, RNA, two $MG^{2+}$ cations and: a UTP molecule, a RTP molecule, a UTP and a Remdesivir nucleotide placed in position i+1 to i+4. RMSD for the different systems during our MD simulations are shown below.

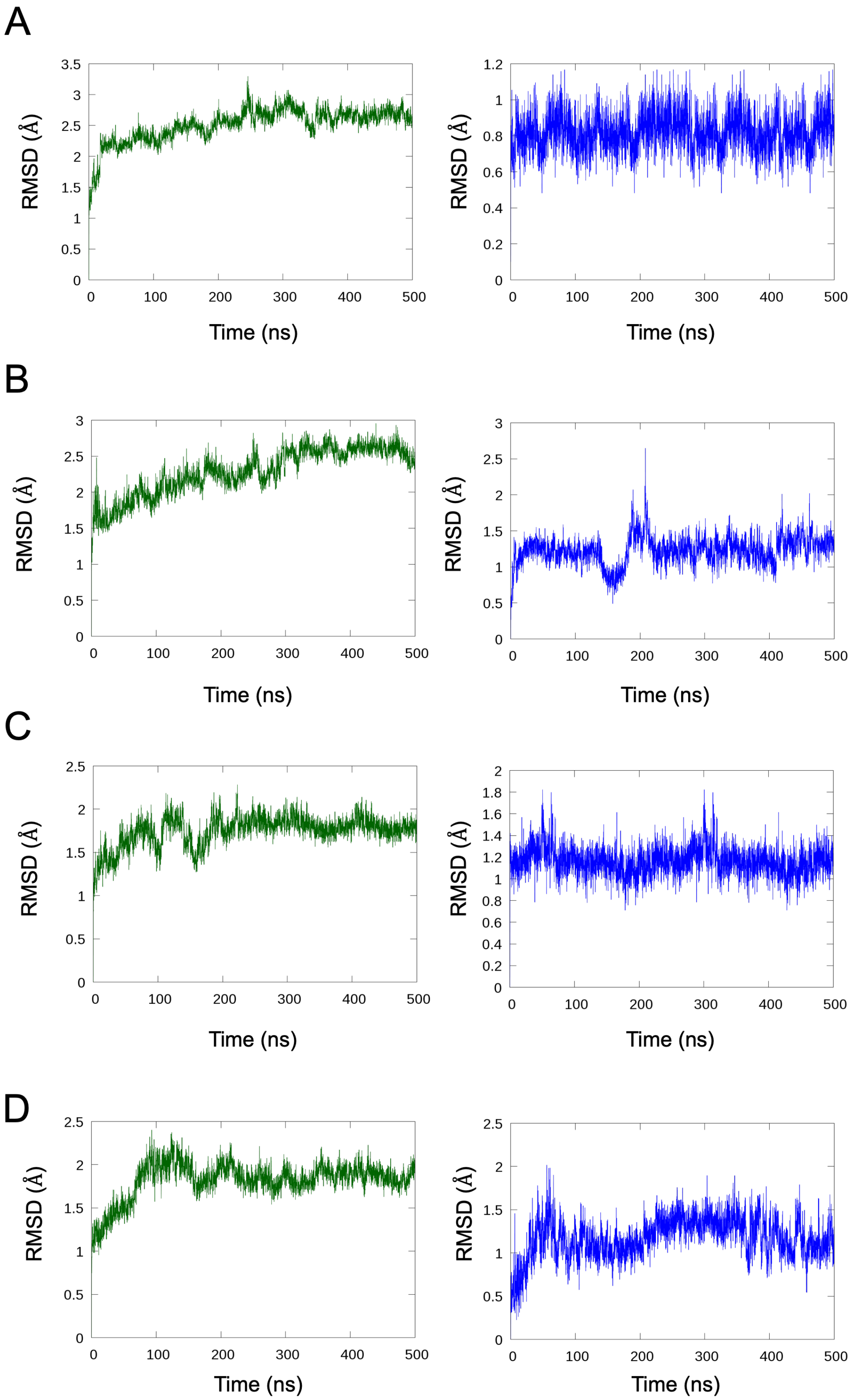
A
B
C
D
RMSD (Å)
Time (ns)
0
100
200
300
400
500

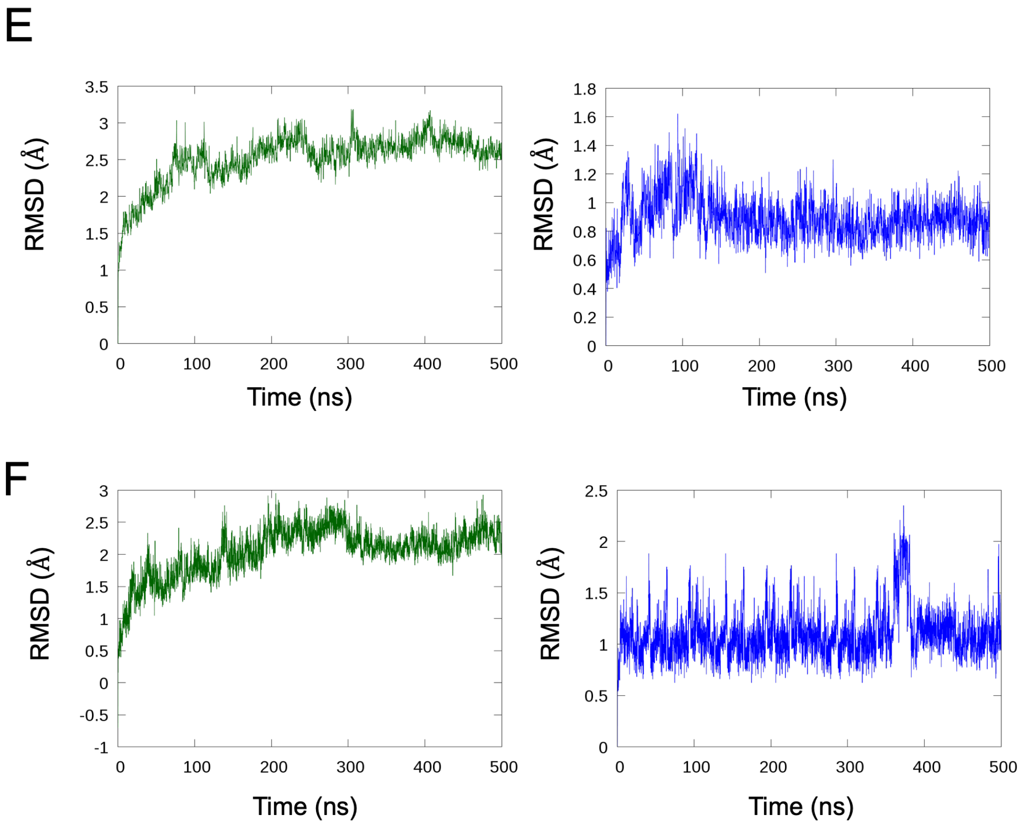


**Supplementary Fig. 15** RMSD for the protein (green) and nucleic acid (blue) backbone atoms during MD simulations. **A** RMSDs for the system containing a UTP molecule. **B** RMSD for the system containing an RTP molecule. **C** RMSD for the system containing a Remedesivir incorporated to the nascent RNA strand. **D** RMSD for the system incorporating one more nucleotide after Remedesivir. **E** RMSD for the system incorporating two more nucleotides after Remedesivir. **F** RMSD for the system incorporating three more nucleotides after Remedesivir.

#### *Structural analysis of Remdesivir inside an RNA double helix*

We performed MD simulations of 1 μs long in two double-stranded RNA dodecamers. One was used as a control a contained only natural-occurring nucleotides with the sequence r(CGCGAAUUGCGC)·r(GCGCAAUUCGCG) while in the other a Remdesivir was placed in the central position r(CGCGARUUGCGC)·r(GCGCAAUUCGCG). The double-stranded RNA molecules where built making use of AMBER Nucleic Acid Builder module. Two A-RNA molecules were constructed. MD simulations were conducted with same protocol and parameters as the ones already explained in the previous section. RNA helical base-pair step parameters were calculated by making use of

CURVES+ and CANAL programs.[17]

### *QM/MM Calculations*

We selected snapshots of the last 50ns as our starting point to build our QM/MM models. The AMBER program making use of the interface with Terachem 1.9 program[18,19] or Gaussian16 program[20] were used. All calculations were performed with electrostatic embedding. For the ligation reaction, the QM subsystem consisted on the UTP or RTP molecule, the terminal nucleotide's sugar ring without the nucleobase, two magnesium ions, and both the side-chain of the protein residues and the waters involved in its coordination sphere (see Supplementary Fig. 4 A). The total number of QM atoms were 117 including the link atoms when a UTP molecule was studied and 122 when a RTP was present. For the covalent adduct formation between the Ser861 and Remdesivir the QM subsystem is depicted in Supplementary Fig. 4 B. The total number of QM atoms including the link atoms were 46 for this step (see Supplementary Fig. 4 B).

We used the link atoms procedure as implemented in the AMBER program to saturate the valence of the frontier between the QM and the MM subsystems. After the system was built the system was re-equilibrated at the QM/MM level by performing minimizations and a 10 ps long NPT QM/MM-MD simulation at the B3LYP/6-31G*/MM level using periodic boundary conditions with an electrostatic cutoff of 12 Å for the QM/MM electrostatic interactions. Afterwards, in order to explore the potential energy surface associated to the reactions we extracted a spherical droplet from the equilibrated structure maintaining. This was done in order to avoid discontinuities in the PES and we simulated an aperiodic system with a cutoff larger than the simulated system. The two-dimensional PESs were calculated at the B3LYP/6-31G**/MM levels, going forward and backward in the reaction process (see Supplementary Fig. 11). The Reaction Coordinates (RC) employed in the exploration of the 2D-PESs of the $Ser_{861}$ covalent addition (see Supplementary Fig. 11) were the simple distance RC1=d1, and the antisymmetric combination of distances d2, d3, d4 and d5 (RC2=[(d2-d3) + (d4-d5)]).

***Exploration of the Minimum Free Energy Paths and Potential of Mean Force***

By means of the string method[21] we investigated the preferred minimum free energy paths (MFEP) by performing QM/MM-MD simulations. We selected snapshots of the last 50ns of the MD simulations as our starting point to build our QM/MM models. The QM subsystems are shown in Supplementary Fig. 4, and atoms were described at the DFTB3[22,23]/MM level, with corrections at B3lyp/6-311++G** to the electronic energy. In Supplementary Fig. 4 is depicted the active space consisting on 2 (d1 and d2 in Supplementary Fig. 4 A) or 5 (d1 to d5 in Supplementary Fig. 4 B) distances that were selected to trace the MFEPs. Afterwards a collective variable was defined along the path[24,25] for a given reaction mechanism and was used to obtain the potential of mean force (PMF) using the umbrella sampling technique.[26] Each MFEP was computed by using 60 string nodes for the phosphoryl transfer reaction and 120 string nodes for the $Ser_{861}$ addition. During the adaptive string optimization the positions and force constants of umbrella sampling windows were taken from the adjusted node parameters.[27] A time step of 1 fs was employed in all cases. Temperature was set to 298K. For the determination of MFEPs the averaged positions of the string nodes were determined in the last 20 ps after the string had converged. Different initial guesses were employed to explore all possible reaction mechanisms. Afterwards 120 points were interpolated for each MFEP between the converged string nodes. These points were used to define the collective variable (s)[24,25] which measures the advance of the system along the MFEP. Umbrella sampling windows were simulated during 20 ps for a relaxation run and during 200 ps during the production run. The time step employed was the same used in the calculation of the MFEP. The statistical uncertainties were calculated as 95% confidence intervals and reached values within ± 1 $kcal \cdot mol^{-1}$ in the free energy. Finally, interpolated corrections[24,25] were made to the DFTB3/MM at the high level B3lyp/6-311++G**/MM in the following way. From the structures collected during the PMF production we performed minimizations in each of the nodes of the MFEP for 1000 minimization steps. Then, single point energy calculations at both the B3lyp/6-311++G**/MM and DFTB3/MM were performed. Finally, the corrections were applied as follows:

$$E = E_{QM}^{LL} + E_{QM/MM}^{LL} + E_{MM} + \mathrm{Spl}[\Delta E_{LL}^{HL}(s)]$$

where Spl is a one-dimensional cubic spline function and its argument, $\Delta E_{LL}^{HL}$ , is the correction term obtained as the difference between the single-point high-level (HL) energy of the QM system and the low level one (LL).
